# Multi-omics discovery of hallmark protein and lipid features of circulating small extracellular vesicles in humans

**DOI:** 10.1101/2024.03.16.585131

**Authors:** Alin Rai, Kevin Huynh, Jonathon Cross, Qi Hui Poh, Haoyun Fang, Bethany Claridge, Thy Duong, Carla Duarte, Jonathan E Shaw, Thomas H Marwick, Peter Meikle, David W Greening

**Author notes:** To whom correspondence should be addressed: Alin Rai, PhD Baker Heart and Diabetes Institute 75 Commercial Road, Melbourne, 3004, Australia David W. Greening, PhD Baker Heart and Diabetes Institute 75 Commercial Road, Melbourne, 3004, Australia.

## Abstract

Extracellular vesicles (EVs) are now being increasingly recognized as an essential signaling entity in human plasma, linking them to health and various diseases. Still, their core protein and lipid componentry, which lie at the center of EV form and function, remain poorly defined. Achieving this unmet milestone remains greatly hindered by abundant non-vesicular extracellular plasma components (non-EVs) in mass spectrometry-based analyses. Here, we performed high-resolution density gradient fractionation of over 140 human plasma samples to isolate circulating EVs, and systematically construct their quantitative proteome (4500 proteins) and lipidome (829 lipids) landscapes. This led to the discovery of a highly conserved panel of 182 proteins (ADAM10, STEAP23, STX7) and 52 lipids (PS, PIPs, Hex2Cer, PAs), providing a deep survey of hallmark molecular features and biological pathways intrinsic to circulating EVs. We also mapped the surfaceome diversity, identifying 151 proteins on the EV surface. We further establish a set of 42 proteins and 114 lipids features that served as hallmark features of non-EV particles in plasma. We submit ADAM10 and PS(36:1) as conserved EV biological markers that precisely differentiate between EV and non-EV particles. Our findings, which can be explored via an open-source Shiny web tool (evmap.shinyapps.io/evmap/) will serve as a valuable repository to the research community for a clearer understanding of circulating EV biology.

## Introduction

Extracellular vesicles (EVs) are membrane-enclosed nanoscale particles (30–1000Lnm in diameter) released by cells into their extracellular space^1,2^. By transferring bioactive cargo such as proteins, lipids, nucleic acids, and metabolites between cells^3–6^, EVs execute diverse biological functions in various physiological and pathological processes. EVs are also found ubiquitously in the human circulatory system^7–15^; with an estimated 50 x 10^6^ particles per ml of plasma^9^, and now recognized as an essential signaling entity in human plasma. While their precise functions in humans remain mostly elusive, circulating EVs have been implicated in essential life processes, such as immune regulation^16^, inter-organ crosstalk^17,18^, tissue homeostasis and regeneration^19^ and coordinating physiological responses^20–22^. Dysregulation of their cargo composition is also implicated in various human diseases such as cancer^23^, cardiovascular disease^24^, artery calcification^20^, and COVID-19 pathogenesis^25^. As minimally invasive liquid biopsies, they have also garnered interest for their potential diagnostic value^9^ and real-time monitoring of therapeutic response^26,27^.

Despite their wide biological and biomedical implications, the study of circulating EVs remains exceptionally challenging^28^, with their conserved protein and lipid componentry remaining poorly characterized and an unmet milestone. Given that conserved molecular blueprint epicentres form and function of a biological system - sharing unique proximity to phenotype and pathophysiology^29^ - closing this knowledge gap is critical for advancing our fundamental understanding of circulating EVs and harnessing their clinical potential. Recently, while several seminal studies^30,31^ have begun to define the precise core of EVs using in vitro culture systems, making such discoveries for EVs circulating in human plasma remains a formidable challenge. This is mainly due to the presence of large abundance of non-EV components (such as lipoprotein particles, soluble proteins, complement proteins, immunoglobulins) in plasma that outnumber EVs by six to seven orders of magnitude^32^. Such non-EV particles co-purify with EVs which invariably limits mass spectrometry-based quantifications primarily to high-abundant plasma proteins or lipoproteins particle-associated lipids, resulting in incomplete and low-coverage data^14,23,25^. Thus, there is an ongoing international effort within the EV community^9,11,31,33–35^ to develop and refine plasma EV isolation methods, with the aim of precisely defining their molecular landscape^10,12,14,36,37^.

A systematic construction of the conserved protein and lipid componentry of circulating EVs in humans has several implications. It will inform us on fundamental building blocks of circulating EVs, providing high-confidence molecular maps and associated biological pathways such as biogenesis (including membrane curvature, stability, cargo recruitment), release, environmental interactions, and uptake mechanisms in humans. Given that existing EV markers obtained from cell culture systems have limited conservation in humans^23^ - potentially due to unique architecture and context of human tissues^38,39^ – these conserved features will also serve as robust EV markers applicable to human plasma that can be rigorously implemented in large-scale population studies, advance EV purification and characterization techniques, and extend international EV guidelines^28^ for standardized circulation EV research. Additionally, these conserved features will bridge the knowledge gap between humans and cultures/animal models, enhancing knowledge transferability and translatability. Other implications include informing on strategies for engineering and functionalizing EV membranes for improved drug delivery vehicles (e.g., extended half-life in circulation), and developing safer EV-based therapies.

In this study, we employed high-resolution density gradient separation to isolate a major EV sub-type, called small EVs from human plasma. We verify the enrichment strategy and EV identity using various biochemical and biophysical characterization, ensuring a high degree of separation of EVs and non-EV particles in plasma. We then construct their detailed proteome and lipidome maps, identifying 182 proteins and 52 lipids as core protein and lipid componentry of EVs, which we refer to as EV hallmark features. We also identify 29 proteins and 114 lipids that are defining features of non-EV particles. These markers, in particular ADAM10 and PS(36:1), enable precise differentiation between EV and non-EV particles using machine-learning. To enhance data accessibility, we developed an open-source R/Shiny web tool (<u>evmap.shinyapps.io/evmap/</u>).

## Results

### Isolation of circulating EVs from human plasma

While there is no method that can isolate EVs to absolute purity, high-resolution iodixanol-based density gradient separation (DGS)^40–42^ remains a powerful strategy for enriching EVs from complex biofluids such as plasma. To this end, we subjected plasma to top-down DGS **(Fig. 1A-B)** which resolved the majority of signals for abundant plasma components such as albumin, apolipoproteins, and argonaut 2 (AGO2, known to associate with non-EV RNA extracellularly^43^) to DGS fractions 1-5. In contrast, CD63 (tetraspanins that are found in small EVs) resolved in DGS fractions 6-8 (corresponding to flotation buoyancy of ∼1.09 g/ml, typical of EVs) **(Fig. 1A-B)**. This resolution remained inaccessible to ultracentrifugation (100,000 *g*, referred to as p100K) alone that co-isolated large quantities of abundant plasma components **(Supplementary** Fig. 1A).

**Figure 1.**
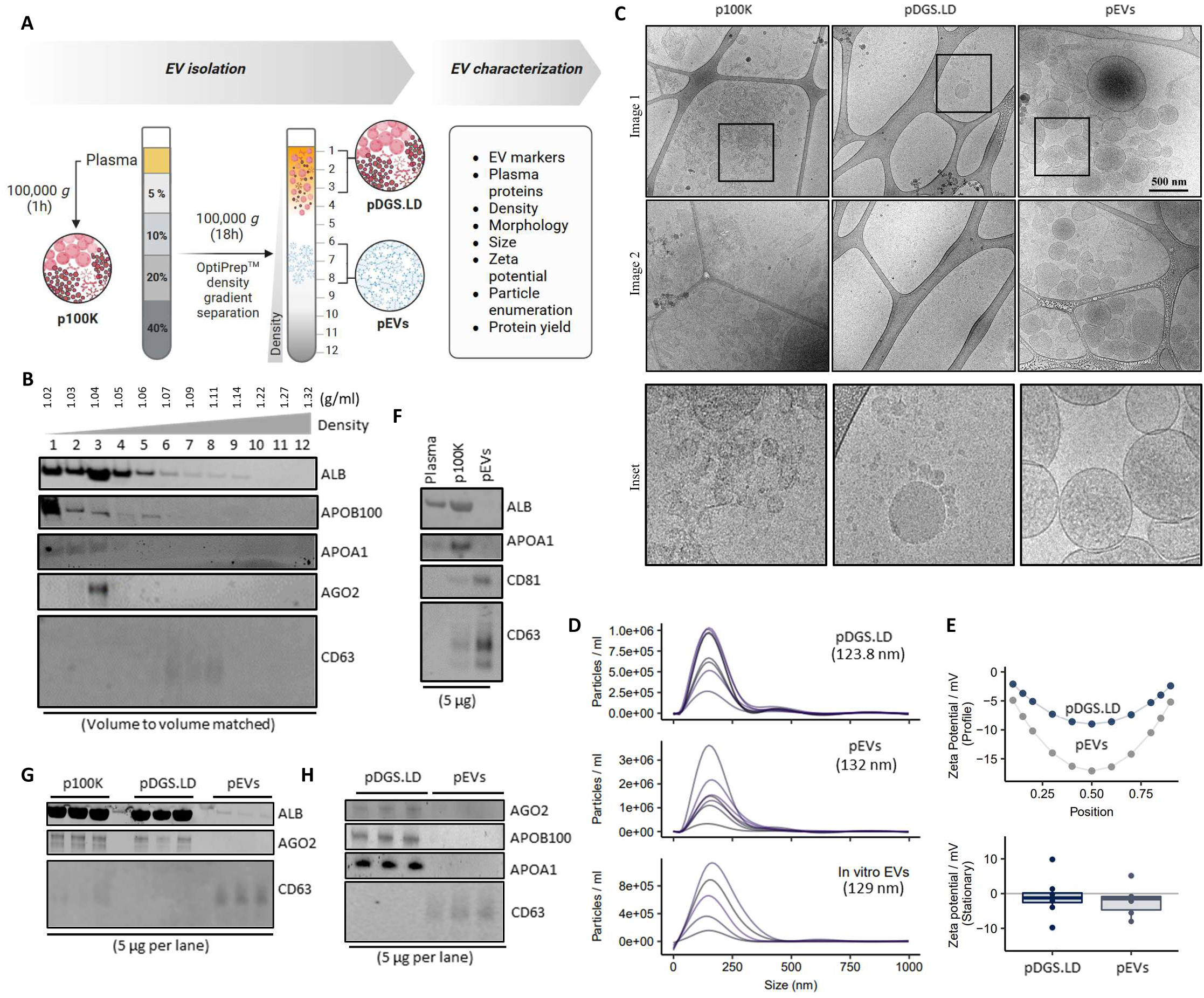
Isolation and characterization of EVs from human plasma. **A.** Workflow for density gradient separation (DGS) of human plasma (0.5 ml) for EV isolation and characterization. **B.** Western blot analysis of twelve DGS fractions (vol:vol matched) of human plasma with antibodies against indicated proteins. **C.** Cryogenic electron microscopic images of p100K, DGS fractions (1-3) and (6-8). Scale bar, 500 nm (n=3). **D.** Size distribution (particle diameter (nm)) of plasma DGS fractions (1-3) and (6-8) (n=7) based on nanoparticle tracking analysis. Small EVs released by SW620 cells (in vitro EVs (n=7)) were also analyzed. **E.** The zeta potential of DGS fractions (1-3) and (6-8) at 11 positions throughout the sample cell (top panel), or stationary layers in the sample cell (bottom panel) (n=5). **F.** Western blot analysis of unprocessed plasma, p100K and DGS fractions 6-8 (pooled). Protein load per lane as indicated. **G-H.** Western blot analysis of indicated DGS fractions from 6 different pooled plasma samples (n=6 independent plasma samples).

EV particles in DGS (6-8) fractions (pEVs; plasma EVs) were morphologically intact, membrane-limited spherical vesicles, consistent with previous reports^44–47^ **(Fig. 1C)**, with a mean diameter of 220.4 nm (**Supplementary** Fig. 1B). In contrast, in p100K contained abundance of proteinaceous material that formed aggregates, whereas DGS (1-3) fractions (referred to as pDGS.LD; plasma DGS light density particles) contained spherical structures without an apparent lipid membrane (resembling size and morphology of lipoprotein particles^8^).

Moreover, nanoparticle tracking analysis **(Fig. 1D)** revealed that pEVs ranged from 30-300 nm in size and displayed a net negative charge **(Fig. 1E)**, consistent with a previous report^48^. Thus, our data show that we can successfully enrich for small EVs from human plasma with minimal contamination from non-EV plasma components. We further demonstrate EV enrichment from multiple plasma samples **(Fig. 1D-E)**. From 1 ml of plasma, we obtained ∼8.7 µg of pEVs (a striking >24,000 fold less compared to total plasma protein) **(Supplementary** Fig. 1C), which enumerated to ∼4.2 x 10^9^ particles **(Supplementary** Fig. 1D-E). To assess separation efficiency, we also performed bottom-loaded DGS by carefully loading plasma at the bottom of the gradient before ultracentrifugation **(Supplementary** Fig. 1F). While EVs were recovered in similar fractions as in the top-loaded approach (CD63 signal in fraction 6), bottom-loaded DGS resulted in greater co-fractionation of non-EV components, including ALB and APOA1, across multiple fractions. This suggests that, compared to top-loaded DGS, bottom-loaded DGS does not achieve the same resolution in plasma samples.

### Constructing the proteome draft of circulating EVs in humans

We next performed MS-based proteomics analysis of pEVs from 38 human plasma samples from multiple sources **(Supplementary** Fig. 2A). To identify EV specific proteins, we compared the pEV proteomes with those of pDGS.LD, p100K and unprocessed plasma, collectively termed as “NonEVs” (n=42) **(Fig. 2A, Supplementary** Fig. 2B**, Supplementary Table 1-6)**. Using stringent peptide and protein identification criterion (1% false discovery rate), we quantified 4631 proteins in pEVs and 1678 in NonEVs **(Supplementary** Fig. 2C**, Supplementary Table 6)**. The size of pEV proteome dataset was comparable to those obtained for in vitro EV proteome data from four different cell lines (4492 proteins), ensuring sufficient proteome coverage.

**Figure 2.**
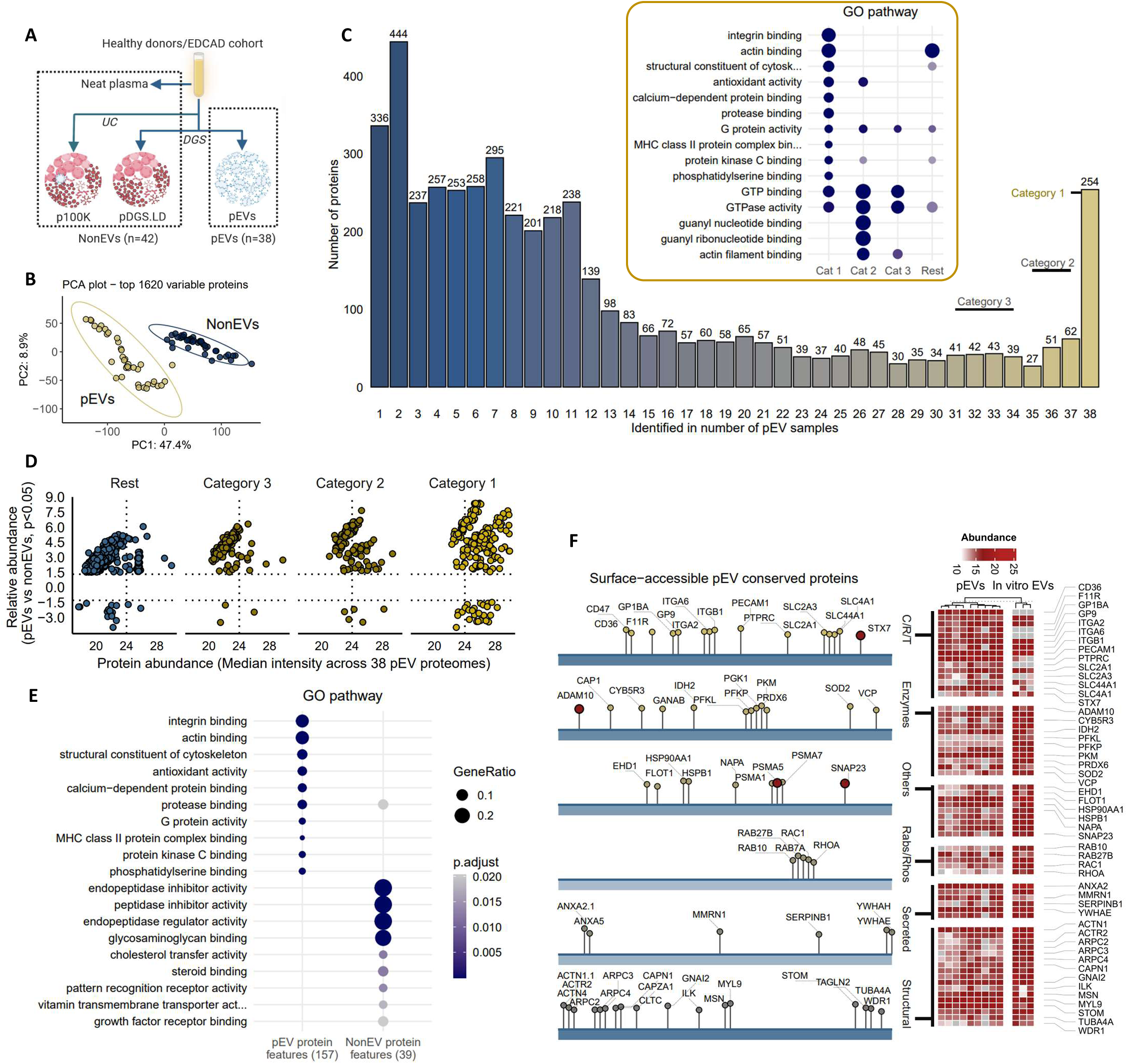
Mapping the core proteome of circulating EVs in humans. **A.** Proteome landscape construction of plasma EVs (pEVs, n=38) and NonEVs (n=42). **B.** Principal component analysis of quantified proteins. **C**. Occurrence analysis of proteins in 38 pEV proteomes where Category 1 proteins represent ubiquitously quantified proteins. Gene Ontology pathways enriched in each category proteins are indicated. **D.** Scatter plot representing differential abundance (p<0.05) of Category 1-3 proteins in pEVs compared to NonEVs. **E.** KEGG pathways enriched (p<0.05) in pEV or NonEV protein features. **F.** Surface-accessible Category 1 proteins of pEVs categorized based on their molecular/functional annotation. C/R/T represents cluster of differentiation (CDs), receptors and transporters. Heatmap for abundance of indicated proteins in pEVs (n=9) and in vitro EVs (n=3).

The pEV proteome displayed a remarkable dynamic range **(Supplementary** Fig. 2D), including extensive coverage of low-abundant circulating proteins^49^ **(Supplementary** Fig. 2E), and was distinct to NonEV proteome **(Fig. 2B, Supplementary** Fig. 2F). Moreover, differentially abundant proteins in pEVs were enriched for terms/pathways associated with small EVs (regulation of actin cytoskeleton and endocytosis) **(Supplementary** Fig. 3A-C**, Supplementary Table 7-8)** and EV biogenesis proteins^30,45,47,50,51^ **(Supplementary** Fig. 3D), while NonEVs contained components related to complement and coagulation, and abundant plasma proteins^52^ **(Supplementary** Fig. 3A-D), further supporting our EV enrichment pipeline at omics level. Importantly, although activated platelet EV proteins such as CLEC1B, PF4 and PPBP (which we previously reported^53^) contribute towards the pEV proteome landscape **(Supplementary** Fig. 4**, Supplementary Table 9)**, we observed an abundance of non-platelet EV proteins (**Extended Data 1A-C)**, suggesting that the pEV proteome represent diverse cellular source^15,54,55^ **(Extended Data 1D-E, Supplementary Table 10)**. Indeed, we quantified diverse cell-, tissue-, and organ-associated proteins in pEVs^56,57^, suggesting that our dataset provides a snapshot of the diverse vesicular population in circulation **(Extended Data 2A-D, Supplementary Table 11)**: many of these cell signatures were also enriched in pEVs vs NonEVs **(Extended Data 2E-F, Supplementary Table 12)**. Furthermore, our high-resolution mass spectrometry quantified low-abundant but biologically significant molecules in pEVs, including signal transduction proteins, cytokines and chemokines, kinases^58^, cell-surface receptors, transporters, and CD proteins^59^, RNA-binding proteins^60^, and transcription factors^61^ **(Extended Data 3A-B, Supplementary Table 11)**. These include RNA binding proteins such as HNRNPK^62^ and PCBP2^63^, TFs such as NME2^64^, kinases such as SRC^65^, chemokines such as TGFB1^66^, and receptors such as integrins^67^, previously reported to be functionally active cargo of EVs. These molecules (e.g., cytokines) exhibit similar abundance to canonical CD proteins typical to EVs, strongly supporting their active incorporation (**Extended Data 3C-D)**.

### Defining the core protein features of circulating EVs

Echoing a recent report^23^, although several EV proteins (namely, 22 core proteins reported in in vitro EVs^30^ and 94 EV marker proteins recommended by the MISEV guidelines^28,36^) were detected in pEVs **(Supplementary** Fig. 5A**, Supplementary Table 6)**, many were not universally quantified **(Supplementary** Fig. 5B-D).

On the other hand, occurrence analysis identified 259 proteins ubiquitously quantified in all 38 pEV samples (termed Category 1 proteins, **Fig. 2C, Supplementary Table 7)**; these proteins were enriched for EV related terms/processes (**Extended Data 4A, Supplementary Table 13)** and encompass proteins associated with endosomal trafficking network **(Extended Data 4B)** such as flotillins, integrins, and CD proteins, whose close interconnectedness is highlighted by their protein-protein interaction network **(Extended Data 4C)**.

For pEV protein feature selection, we first selected proteins that were 100% conserved in pEV proteomes (i.e., present in all 38 proteome datasets, Category 1 proteins), which resulted in 259 proteins. Next, of these 259 proteins, we selected proteins with fold change >1.5 and p-val < 0.0001 in pEV vs. NonEVs, resulting in 182 proteins, which we refer to as ‘pEV protein features’ (**Fig. 2D, Extended Data 5, Supplementary Table 7**, FDR <0.05 as reported in **Supplementary Table 20**).

For NonEV protein feature selection, we first selected proteins that were 100% conserved in NonEV proteomes (i.e., present in all 42 proteome datasets), which resulted in 114 proteins. Next, of these 114 proteins, we selected proteins with fold change < -1.5 and p-val < 0.0001 in pEV vs. NonEVs, resulting in 42 proteins, which we refer to as ‘NonEV protein features’ (**Extended Data 6, Supplementary Table 7**, FDR <0.05 as reported in **Supplementary Table 20**).

Emphasizing their significance as essential EV components, the pEV protein features were enriched for classical EV pathways **(Fig. 2E, Supplementary Table 14)**. These pEV protein features showed coordinated molecular patterns across functional groups associated with “vesicular transport”, “actin cytoskeleton regulation”, and “membrane-raft assembly” (see **Extended Data**D**7A** for enrichment-based associations). Upon closer inspection, these correlated molecular patterns were conserved across individual EV proteomes at the cellular level or within specific EV subpopulations (i.e., CD9/CD81/CD63-positive EVs) (**Extended Data**D**7B, Supplementary Table**D**15**). Moreover, these enrichment trends were observed in EVs from primary human fibroblasts (**Extended Data**D**7C**) and were enriched in EVs relative to parental cell proteomes^30^ (**Extended Data**D**7D**). In contrast, NonEV protein features were enriched for “endopeptidase inhibitor activity, “Complement and coagulation cascades”, and “Cholesterol metabolism” (**Fig. 2E, Supplementary Table 14)**.

We next queried whether the pEV protein features also comprised the EV surfaceome, essential interactive surface platforms on EVs^68^. To explore this, surface-accessible proteins on pEV were labelled with membrane-impermeant Sulfo-NHS-SS-Biotin, captured with avidin-coated beads, and analysed through MS-based proteome profiling (n=9) (**Supplementary** Fig. 6A**, Supplementary Table 16)**. Our data revealed that 151 pEV protein features were surface-accessible, including CD proteins (CD44, CD47), integrins (ITGA2, ITGA6, ITGB1) and annexins (ANXA2, ANXA5) **(Fig. 2F, Supplementary** Fig. 6B-C). We further confirmed their surface-accessibility in in vitro EVs **(Fig. 2F, Supplementary Table 16)**. In contrast, the conserved SDCBP protein remained inaccessible to biotin-capture, which is in direct agreement to its luminal localization^30^.

By leveraging data from published proteomes^15,69,70^, we demonstrated a remarkable conservation of pEV protein features in multiple plasma or serum EVs **(Fig. 3**, panel **A-C, Supplementary Table 7)**. These features were also conserved in different EV sub-populations (CD81+, CD63+, and CD9+ EVs) prevalent in the circulatory system **(Fig. 3**, panel **D-F)**, with 100% quantification of at least 74 proteins. Moreover, interrogating previous published sEV proteomes from fourteen cell lines^30^, 81 EV features were also conserved, displaying greater abundance in EVs compared to cells (**Supplementary** Fig. 7A**-B)**. Additionally, we experimentally verified their conservation in in vitro EVs (n=12) release by four cell lines **(Fig. 3, I-L)**. Finally, we validated their conservation in pEVs (n=4) and in vitro EVs (n=6) using orthogonal TMT-based isobaric multiplexed proteomics **(Fig. 3**, panel **G-H, Supplementary Table 26)**. Overall, 78 pEV proteins displayed 100% conservation, of which 63 were surface-accessible (**Supplementary Table 7)**.

**Figure 3.**
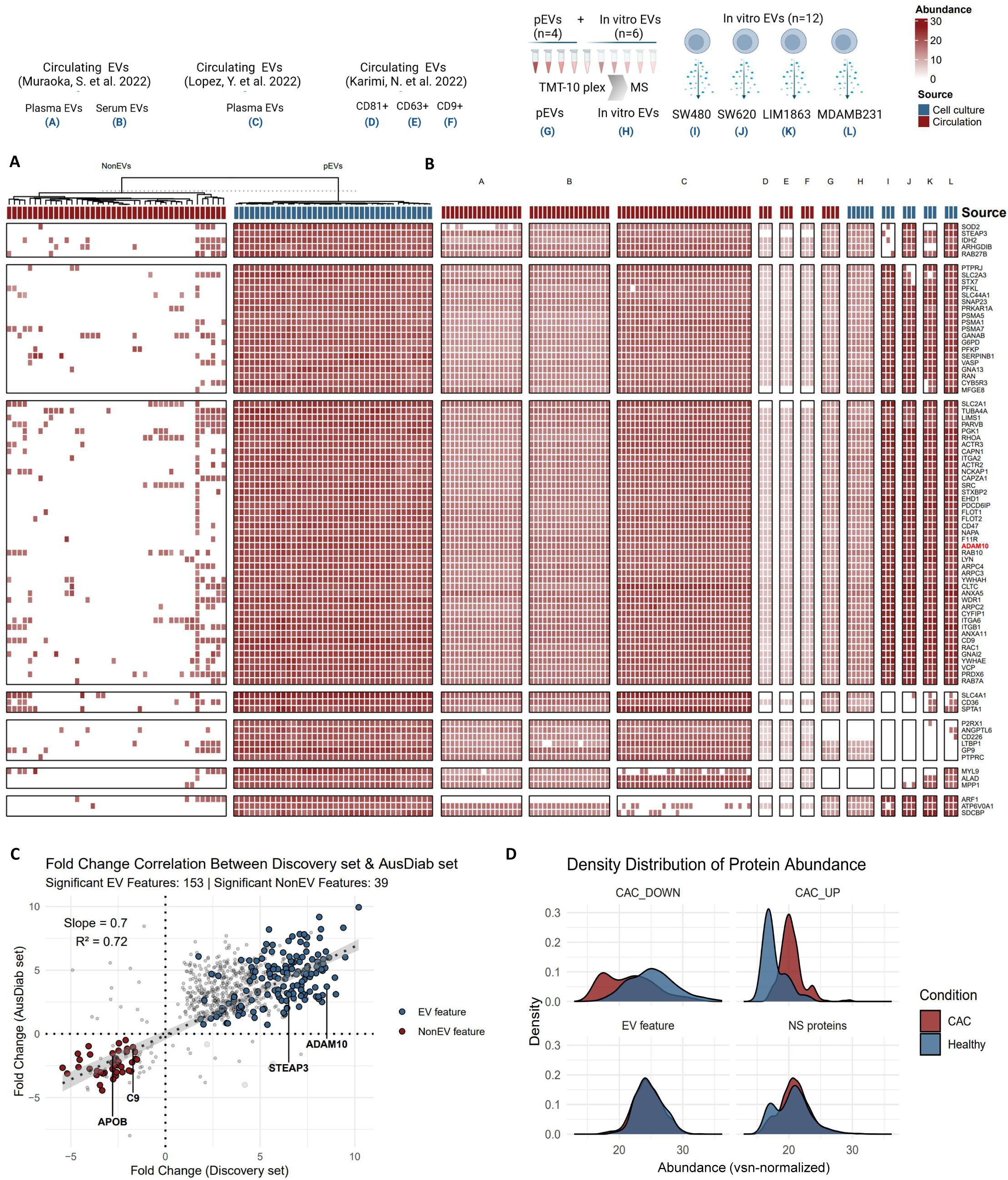
Conservation of circulating EV protein features. **A.** Heatmap depicts quantification of EV protein features in our pEV and NonEV proteome datasets (only protein features quantified in <30% of NonEV datasets were used). **B.** Heatmap depicts quantification of EV protein features in previous reported proteomes for circulating EVs: (A-C) EV preparations from plasma or serum sourced from healthy individuals or those with known pathology^15,69^, and (D-F) EV subtypes (CD81+, CD63+, and CD9+ EVs) prevalent in the circulatory system^70^. As an orthogonal validation, heatmap also depicts conservation of EV protein features (G-J) using TMT-based isobaric multiplexing of pEVs (n=4) and in vitro EV proteomes (n=6), and (I-L) using label-free quantitative proteomics for in vitro EVs (n=3 per cell line). **C.** Scatter plot of fold change (FDR < 0.05) EV protein features and NonEV protein features in discovery set and validation set (AusDiab set). Rest of the proteins are indicated with grey points. Pearson r = 0.851, p < 2 × 10L¹L. **D.** Density plot of proteins in pEV proteomes from individuals with either positive CAC score (CAC group) or zero CAC score (Healthy group). CAC_DOWN: proteins significantly (FDR <0.05) downregulated in pEVs from CAC vs Healthy. CAC_UP: proteins significantly (FDR <0.05) upregulated in pEVs from CAC vs Healthy. EV feature: 182 pEV protein features. NS_proteins: proteins with similar abundance between pEVs from CAC vs Healthy.

To validate our identified pEV markers, we analysed plasma from an independent external cohort (AusDiab, n=12 pEVs, n=12 NonEVs) using the same EV isolation and proteomic pipeline (**Supplementary Table 17**). Comparative proteomic analysis revealed that 177 out of 182 pEV protein features and all 42 NonEV protein features were detected, with 87.4% (156/182) of pEV proteins and 92.9% (39/42) of NonEV proteins significantly enriched in their respective fractions (FDR < 0.05) (**Supplementary Table 18, Extended Data 8)**, with similar enrichment in Gene Ontology pathways in our discovery cohort (**Supplementary Table 19**).

Comparative analysis of fold change of pEV and NonEV protein features between the discovery and validation datasets (**Supplementary Table 20**) demonstrated a strong positive correlation (Pearson r = 0.851, p < 2 × 10L¹L), confirming the reproducibility of pEV protein features across independent populations (**Figure 3C)**.

Importantly, we show that pEV protein cargo differs between individuals with and without coronary artery calcium (CAC) deposits in our EDCAD cohort **(Extended Data 9A-C, Supplementary Table 21-22)**. Occurrence and development of calcification is a complex biological process, which is regulated by multiple factors, including EVs that are regarded as nidus for calcification by providing mineral nucleation sites^71^. CAC scoring, a predictor of future cardiovascular events, is determined by computed tomography. We identified 76 upregulated and 127 downregulated proteins (FDR < 0.05) in pEVs from individuals with positive CAC scores **(Extended Data 9D-E, Supplementary Table 21-22)**, with the upregulated proteins significantly enriched in the GO pathway ‘Abnormal cardiovascular system physiology’ **(Extended Data 9D, Supplementary Table 23)**. Notably, this includes (Cystatin C (CST3)^72^, TPM2^73^, and TPM1^74^, and TXNRD2^75^, all of which have well-established roles in cardiovascular disease and vascular calcification. Despite these proteomic differences between CAC and non-CAC pEV proteomes, the core set of 182 pEV marker proteins remained highly conserved across both groups **(Extended Data 9F-G, Supplementary Table 24)**, indicating that the fundamental pEV molecular signature is stable across individuals, potentially even in the context of disease (**Figure 3D**). This preservation of core molecular identity in pEVs across CAC and non-CAC individuals, while simultaneously capturing disease-specific molecular shifts in the EDCAD study, reinforces their potential as a well-defined reference for plasma EV research including disease biomarker discovery. In contrast, NonEV particles and neat plasma proteomes were unable to distinguish CAC from non-CAC individuals, underscoring the power of highly purified pEVs (separated from NonEV particles) in revealing disease-associated changes in the plasma proteome **(Supplementary** Figure 8).

To assess the broader biological relevance of pEV protein features beyond plasma-derived EVs, we analysed EV proteomes from non-transformed human fibroblasts and endothelial cells (**Extended Data 10, Supplementary Table 25)**. Among the 182 pEV protein features, 135 were detected in this dataset, with 43 proteins showing 100% conservation across fibroblast- and endothelial-derived EVs. Conserved proteins included ADAM10, integrins, Rabs, Annexin A5, CD markers (CD9, CD44), and SDCBP, reinforcing their role as core EV components across different biological sources. Proteins absent from non-transformed cell-derived EVs primarily included immunoglobulins and complement proteins, supporting their plasma-specific nature or association with the EV protein corona.

Thus, our study defines highly conserved protein features of circulating EVs in human plasma (**Table 1** highlights top 25 EV or NonEV protein features, “*” indicates surface accessible proteins).

### The lipidome draft of circulating EVs in humans

Next, we defined the lipidome landscape of circulating EVs. The majority of membrane lipids fall under glycosphingolipids, sphingolipids and sterols (predominantly cholesterol in mammals). Therefore, to capture this lipid diversity, we employed a high-throughput targeted-lipidomics platform interrogating 829 lipids representing 40 lipid classes within these three major groups **(Fig. 4A, Supplementary** Fig. 9**, Supplementary Table 27)**. Because the lipidome of small EVs from cells are not as well defined compared to their proteome counterpart, we also performed lipidome analysis of in vitro EVs and parental cells using the same platform **(Supplementary** Fig. 9A-B**, Supplementary Table 27)**. We reasoned that circulating EV lipid features should ideally be enriched in in vitro EVs compared to cells.

**Figure 4.**
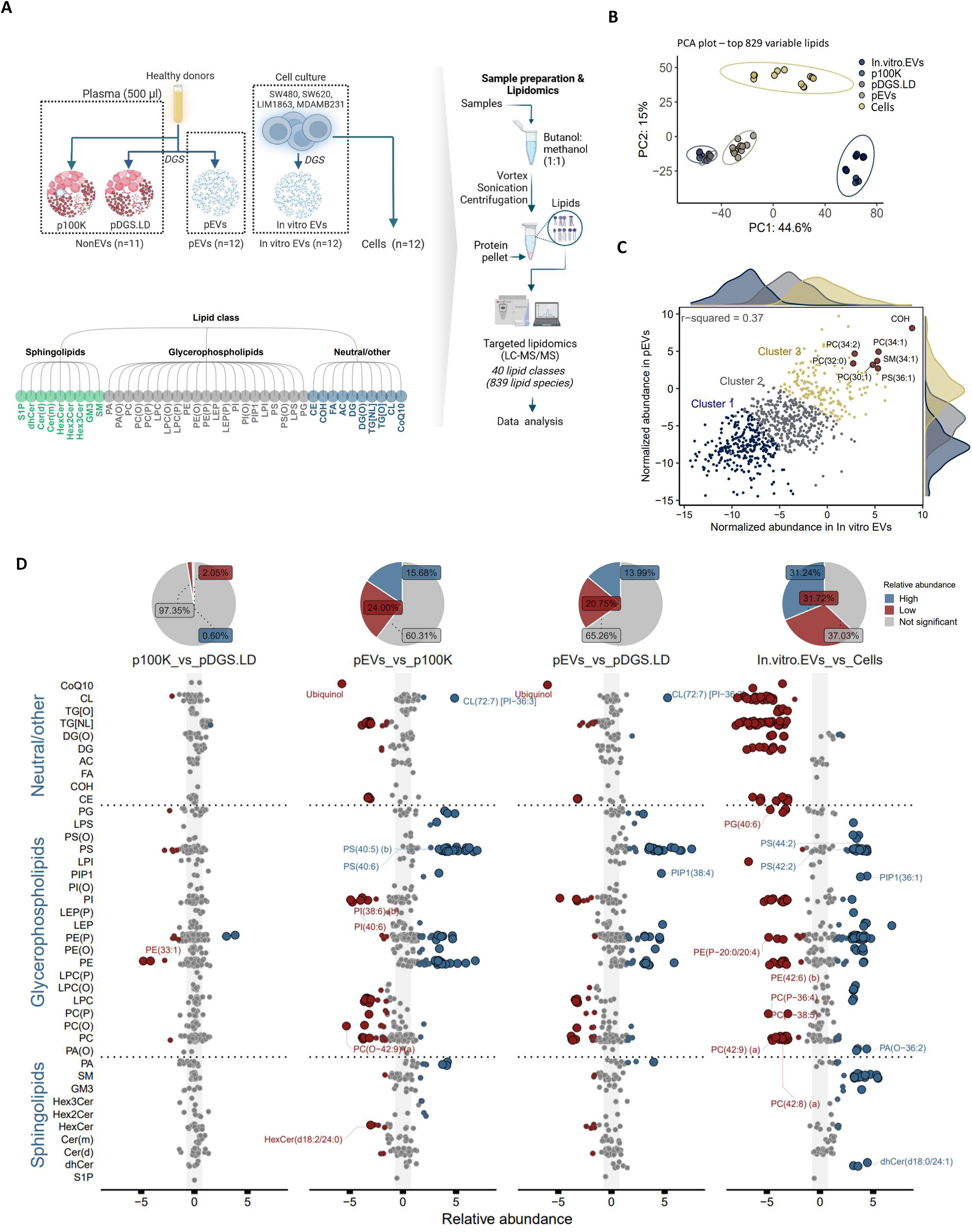
Lipidome landscape of circulating EVs. **A.** Workflow for lipidomic analysis, interrogating 829 lipids representing 40 lipid classes within three major groups. **B.** Principal-component analysis of lipidome data showing group clustering and separation of lipidomes (pEVs; n=12, p100K; n= 6, pDGS.LD; n=5, in vitro EVs; n=12, cells; n=12). **C.** Scatter plot depicts abundance of 829 lipids in pEVs and in vitro EVs with lipids belonging to cluster 3 displaying high abundance in both pEVs and in vitro EVs. **D.** Scatter plot showing relative abundance of lipids in pEVs vs p100K/pDGS.LD and in vitro EVs vs cell lipidome data sets. Blue circles (lipid markers) represent lipids with significantly greater abundance (fold change >1.5 & p<0.05). Red circles (exclusion lipids) represent lipids with significantly lower abundance (fold change <-1.5 & p<0.05). Grey circles represent lipids that do not meet the above criteria. Pie chart depicts number of lipids (%) with differential abundance between the data sets.

Principal Component Analysis revealed that pEVs and NonEVs particles (p100K/pDGS.LD) lipidomes were distinct **(Fig. 4B)**. Highly-abundant lipids in both pEVs and in vitro EVs included cholesterol (COH), phosphatidylcholines (PC), phosphatidylethanolamine (PE), and sphingomyelin (SM) **(Fig. 4C)**; lipids that are well-recognized major structural components in eukaryotic cell membranes^76^ with housekeeping structural functions. For example, PC forms a planar bilayer with COH, ensuring membrane stability and integrity, while the incorporation of conical PE and SM imposes a curvature stress crucial for membrane budding, fission and fusion. These findings align with previous reports highlighting COH as one of the most abundant EV lipids (40–60% of total lipids)^77^.

Over 30% of measured lipid species displayed differential abundance (adjusted p values <0.05, fold change >1.5) between pEVs and NonEVs (p100K/pDGS.LD) **(Fig. 4D, Supplementary** Fig. 10A-B**, Supplementary Table 28)**. Lipids enriched in pEVs are linked to EV biogenesis^77^ and include dihydroceramides (dhCer)^78^, trihexosylceramides (Hex3Cer), dihexosylceramides (Hex2Cer) (**Supplementary** Fig. 10C). These lipids are similarly enriched in in vitro EVs *versus* their parental cells **(Fig. 4D, Supplementary** Fig. 10D-E**, Supplementary Table 28)**. In contrast, lipids enriched in NonEVs include the lipid classes cholesterol ester (CE), triacylglycerol (TG) and coenzyme Q10 (CoQ10)^79^, major components of lipoprotein particles. This further supports our EV enrichment pipeline at lipidomics level.

### The core lipidome of circulating EVs

To identify pEV lipid features, we performed K-means clustering of differentially abundant lipids between pEVs *versus* NonEVs and in vitro EVs *versus* cells **(Fig. 5A, Supplementary Tables 29-30)** which resulted in four major clusters. Representing pEV lipid features were Cluster c4 lipids (52 lipids) enriched in pEVs (compared to NonEVs) and in vitro EVs (compared to cells). In contrast, representing NonEV lipid features were Cluster c1 lipids (114 lipids) enriched in NonEVs and cells.

**Figure 5.**
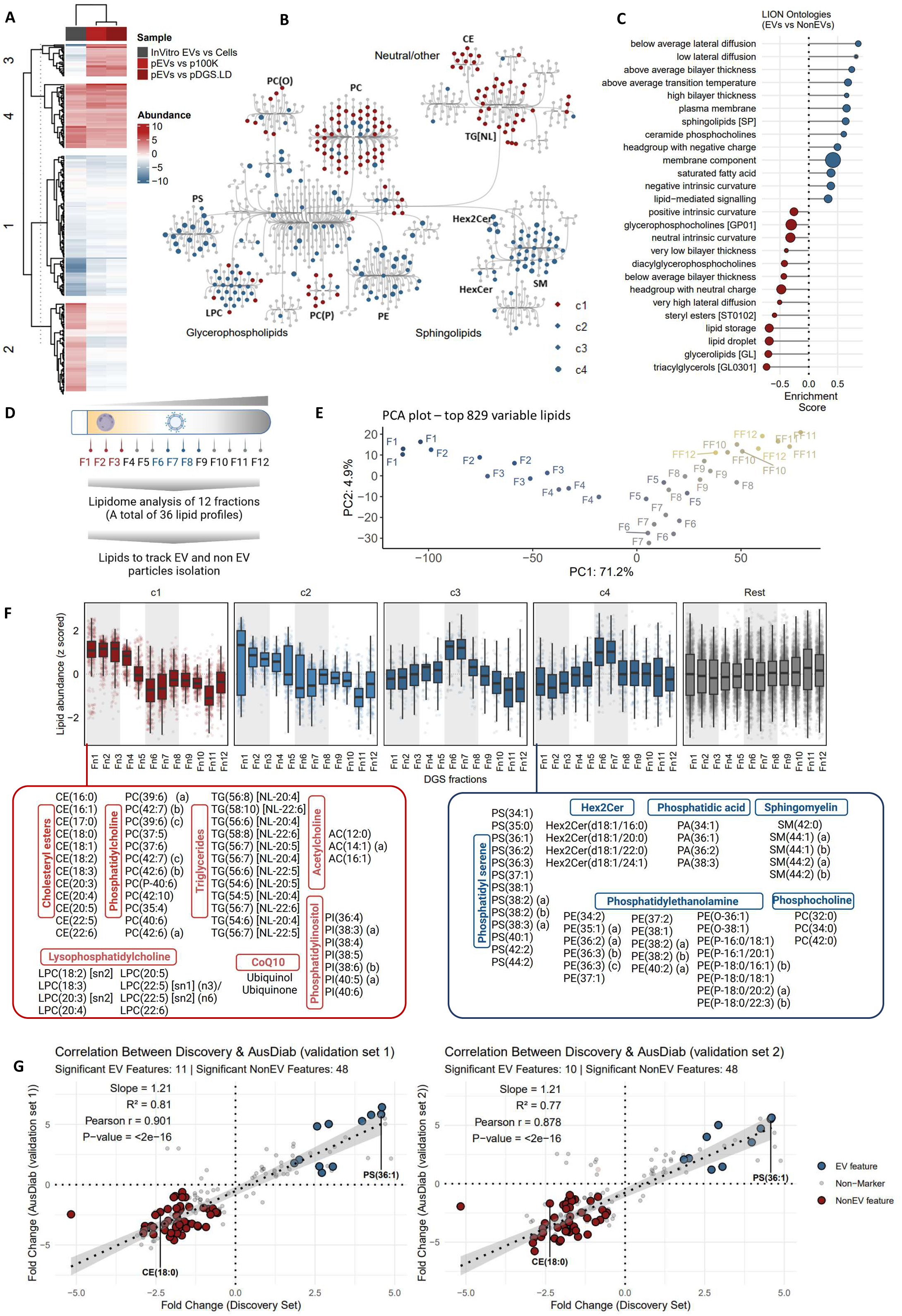
Conserved lipid features of circulating EVs in humans. **A.** Heatmap depicting K-means clustering of differentially abundant lipids from Figure 4D. **B.** Network map of lipids grouped based on lipid classes. Blue circles represent EV-associated lipids clusters c2, c3 and c4, whereas red circles represent NonEV-associated lipid cluster c1. Grey circles are lipids with similar abundance between EVs and NonEVs. **C.** LION Ontologies enriched in EV lipid features (cluster c4 lipids) or NonEV lipid features (cluster c1 lipids). **D.** Density gradient separation of plasma. The twelve fractions were subjected to lipidomics analysis. **E.** Principal component analysis of lipidomes of 12 fractions (n=3). **F.** Box plots depict abundance of cluster 1-4 lipids for DGS fractions. Y axis represents Z-scored abundance (MS-based abundance for each lipid - mean abundance) / standard deviation (Z score normalization). Grey lines mark either fractions 6-7 (corresponding to pEV fractions) or fractions 1-3 (corresponding to pDGS.LD fractions). Lipids from cluster c4 and cluster c1 are depicted. **G.** Scatter plot of fold change (FDR < 0.05) EV lipid features and NonEV lipid features in discovery set and two validation sets (AusDiab set). Rest of the lipids are indicated with grey points.

Notably, we observed a class-level co-regulation of pEV lipid features **(Fig. 5B),** supporting previous reports that lipid species from the same pathway correlated/co-regulated^80^. For example, lipids from PS and PE class were co-enriched in EVs, which is indicative of their co-localization to external leaflets of EV membranes and co-regulating similar cellular processes^81,82^. Another example includes SM and ceramides co-enriched in EVs which is indicative of SM-ceremide axis driven EV biogenesis^78,83^.

Emphasizing their significance as essential EV lipid components, Cluster c4 lipid features **(Extended Data 11)** were linked to LION Ontology terms such as bilayer membrane and plasma membrane **(Fig. 5C)**, but more specifically with EV features such as glycerophosphatidylserine (known to decorate the outer membrane/leaflet of EVs) and low lateral diffusion (a distinct physical property of lipid rafts^84^ that are hubs for EV formation). Conversely, Cluster c1 NonEV lipid features **(Extended Data 11)** were linked to lipid storage/droplets, typical of lipoprotein particles.

To validate the association of Cluster c4 lipids with circulating EVs, and Cluster c1 with NonEV particles in plasma, we subjected twelve fractions of plasma DGS (n=3 plasma samples) to the same targeted lipidomics platform **(Fig. 5D-E, Supplementary** Fig. 11A-B**, Supplementary Table 30-31)**. The normalized abundance of lipids across the fractions were plotted (z-scored within each fraction) **(Fig. 5F)**. Indeed, Cluster c4 lipids were enriched in DGS fractions 6 and 7, where pEVs resolve. In contrast, Cluster c1 lipids were enriched in DGS fractions 1-4, where the majority of NonEV particles resolve, reinforcing the specificity of Cluster c4 lipids for circulating EVs in plasma.

To further assess the conservation of pEV lipid features, we performed lipidomic profiling of pEVs and NonEVs isolated from two sets of plasma from AusDiab study (n=10 plasma samples for validation set 1 & n=12 plasma samples for validation set 2) and non-transformed (primary human fibroblasts and endothelial cells) cell-derived EVs (**Supplementary Table 32, Extended Data 12, Supplementary** Fig. 12). In this validation lipidomic workflow, we re-quantified 12 out of 52 pEV lipid features and 48 out of 114 NonEV lipid features and report their differential abundance analysis (**Supplementary Table 32-33, Supplementary** Fig. 13). Comparison of fold changes between these lipid features across the discovery and validation datasets showed a strong correlation (**Figure 5G**, **Supplementary Table 34,** Pearson r = 0.901, p < 2 × 10L¹L for validation set 1, and Pearson r = 0.878, p < 2 × 10L¹L for validation set 2), confirming the reproducibility of pEV lipid features. The relative abundance of these features was also conserved in EVs from primary human fibroblasts and endothelial cells (**Extended Data 12, Supplementary** Fig. 13). These findings highlight the robustness of our identified EV lipid markers across independent populations and multiple EV sources.

Thus, our study defines highly conserved lipid features of circulating EVs in human plasma (**Table 2** “*” indicates top 25 enriched lipids in either plasma EVs or NonEVs, based on relative fold change in abundance).

### Biological protein and lipid markers for circulating EVs

We next investigated the ability of the EV and NonEV protein feature panel to distinguish between pEVs vs NonEV particles using machine learning (Naïve Bayes algorithm). For this, pEV and NonEV proteomes were evenly partitioned based on sample type into training set (70% of the samples) and a validation set (remaining 30% of samples). By bootstrapping resampling method employed with 25 resampling iterations, the model achieved absolute accuracy (**Supplementary** Fig. 14A). Our model also achieved excellent accuracy (97%) in EV particle identification in an independent test set comprising of pEVs and NonEVs proteomes from additional 16 plasma samples (**Supplementary** Fig. 14A). Moreover, a panel comprising 151 surface accessible EV protein features were also able to distinguish between EV and NonEV particles with 100% accuracy (**Supplementary** Fig. 14B).

To facilitate routine implementation and translational feasibility, we applied Recursive Feature Elimination (RFE) to systematically reduce dimensionality and identify a robust minimal signature with classification accuracy. Of the 182 EV protein features, RFE based random forest algorithm identified ADAM10 as a prominent feature contributing to model performance **(Supplementary Table 35)**. Moreover, ADAM10 protein showed exclusive and absolute quantification in pEVs as well as in vitro EVs, compared to NonEVs (**Fig. 6A**). Moreover, within each EV proteome the abundance of ADAM10 was up to 7.5-fold higher compared to the median intensity (**Fig. 6B, Supplementary Table 36**); this relative abundance of ADAM10 within each proteome alone can distinguish between pEVs vs NonEV particles (**Supplementary** Fig. 14C).

**Figure 6.**
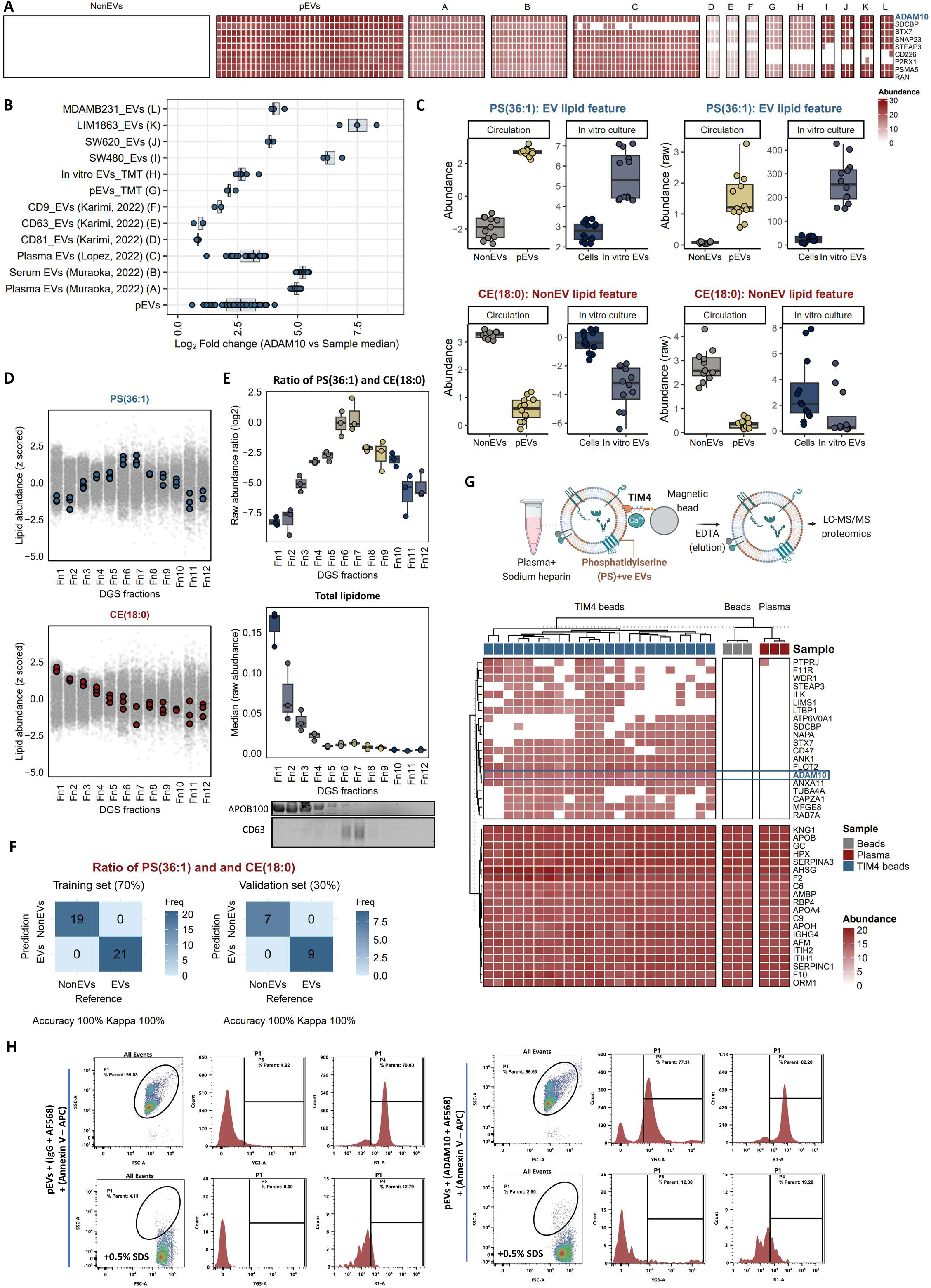
ADAM10 protein and phosphatidylserine lipid are biological markers for circulating EVs in humans. **A.** Heatmap depicting pEV protein features that are uniquely identified in pEVs compared to NonEVs and their conservation, in particular for ADAM10, in previously published circulating EV proteomes^15,69^ (A-C) and (D-F) EV subtypes (CD81+, CD63+, and CD9+ EVs) in plasma^70^, and in our TMT-based isobaric multiplexing of pEVs (n=4) and in vitro EVs proteomes (n=6) as well as label-free proteomes of (I-L) small EVs from 4 cell lines (n=3 per cell line). Ratio of ADAM10 intensity and corresponding sample median intensity for each proteome is depicted in **B** as a boxplot. **C** Box plot depicting normalised intensities (abundance log2-tranformed) and raw intensities of PS(36:1) and CE(18:0) in indicated lipidome datasets. **D.** Abundance of PS(36:1) and CE(18:0) in lipidome datasets across 12 DGS fractions (n=3). Grey points represent rest of the quantified lipids. **E.** Top panel: Relative abundance of PS(36:1) and CE(18:0) in lipidome datasets across 12 DGS fractions (n=3). Middle panel: Median of lipidome datasets across 12 DGS fractions. Bottom panel: Western blot analysis of 12 fractions for indicated proteins. **F.** Confusion matrix using Neural Network algorithm (’nnet’) classifier of training set (70%) and validation set (30%) using relative abundance of PS(36:1) versus CE(18:0). **G.** Workflow for capturing PS+ EVs from human plasma using TIM4-magnetic bead conjugate. Captured EVs (n=23), along with mock capture using beads alone (n=3) or unprocessed plasma (n=3), were subjected to proteomics analysis. Heatmap depicts absolute and exclusive quantification of ADAM10 protein in captured EVs. **H.** Scatter plot showing detection of ADAM10-positive and PS positive pEVs, which was sensitive to SDS detergent (0.5%) solubilization.

Similarly, using a Neural Network algorithm (’nnet’) for machine learning, a panel comprising of EV and NonEV lipid features were also able to distinguish between EVs (pEVs or in vitro EV lipidomes) vs NonEVs (p100K, pDGS.LD or cell lipidomes) (**Supplementary** Fig. 15A). Moreover, this model achieved 97% accuracy in classifying EV particles across DGS plasma fractions (**Supplementary** Fig. 15B).

To enhance translational feasibility, we applied RFE to reduce dimensionality and identify a minimal lipid signature with high classification accuracy. This approach identified PS(36:1) (EV lipid feature) and CE(18:0) as prominent lipid features **(Supplementary Table 35)**, highlighting their marker potential.

Indeed, PS(36:1) lipid is one of the most abundant lipids present in both pEVs and in vitro EVs **(Fig. 4C**), making them amenable to robust and reliable measurements in diverse analytic tools. Moreover, PS(36:1) and CE(18:0) lipid abundances, and importantly their relative abundances, concur with EV protein CD63 and lipoprotein APOB100 resolution on plasma DGS fractions **(Fig 6. D-E)**. Our machine learning (’nnet’) model, based on PS(36:1) and CE(18:0) relative abundance within each lipidome serving as a feature **(Supplementary Table 37)**, can distinguish EVs (pEVs or in vitro EV, DGS fractions 6-7 lipidomes) vs NonEVs (p100K, pDGS.LD or DGS fractions 1-5 lipidomes) with absolute accuracy (**Fig. 6F)**. Because EV and NonEV enrichments along 12 fractions represent a continuum, which can be inferred from PS(36:1) and CE(18:0) relative abundances, our data suggests that these lipids can thus serve as a marker for assessment of EV purity. Importantly, quantitative measurement would be possible in subsequent studies due to the availability of isotope labelled internal standards for these two species, which assist in translation across different laboratories and instrumental setups.

We next investigated whether PS and ADAM10 constitute protein and lipid biological marker componentry on circulating EVs. For this, we employed previously reported^85^ strategy of using extracellular region of T-cell immunoglobulin domain and mucin domain-containing protein 4 (Tim4) immobilized on magnetic beads to directly capture (in the presence of calcium ions) PS+ EVs from plasma (n=23), which was subsequently released (by adding EDTA) and subjected to MS proteomics **(Fig. 6G, Supplementary Table 38)**. Compared to mock capture (beads alone) or unprocessed plasma, PS+ EV proteomes displayed 100% quantification of ADAM10 across all twenty-three samples **(Fig. 6G)**. Additional EV protein features (e.g., FLOT2, RAB7A) were also conserved in these proteomes, which support their EV identity.

Furthermore, single-vesicle analysis using Cytek Aurora flow cytometry confirmed the presence of ADAM10 on a subset of plasma EVs (**Figure 6H**, **Supplementary** Figure 16), with ∼40% of pEVs exhibiting ADAM10^+^ signals, comparable to PS signal detected using Annexin V staining (**Figure 6H**, **Supplementary** Figure 16C**, Supplementary Table 39**). SDS detergent treatment showed strong reduction in fluorescence signal intensity and count, suggesting their EV-origin^86^ (**Supplementary** Figure 16B-C). These findings suggest that while ADAM10 is a component of plasma EVs across individuals, it is not universally present across all vesicles, consistent with previous report^87^.

In our validation set, PS(36:1) lipid was significantly enriched in pEVs and in vitro EVs, reinforcing its role as a conserved pEV lipid marker **(Extended Data 13)**. In contrast, CE(18:0) lipid was enriched in NonEVs and neat plasma. These findings confirm that pEV protein and lipid markers are conserved beyond plasma-derived EVs, reinforcing their biological significance and potential as robust EV classification markers.

Thus, our data show that ADAM10 and the ratio combination of PS(36:1) and CE(18:0) serves as highly conserved and reliable biological markers for EVs in human plasma.

### R/Shiny web tool for EV proteome and lipidome data

Lastly, to facilitate easy access to our data and enhance re-use, we have developed an open-source R/Shiny web tool (<u>evmap.shinyapps.io/evmap/</u>). This tool allows users to quickly interrogate our proteome **(Supplementary** Fig. 17) and lipidome datasets **(Supplementary** Fig. 18) for their molecule(s) of interest. This tool also allows assessment of feature conservation in published studies, surface-accessibility of EV proteins, and construct network analysis, retrieve Gene Ontology and KEGG pathways for selected protein features. Lipid features can also be quickly interrogated for their abundance in EVs vs NonEV particles and their distribution in plasma DGS lipidome fractions. We anticipate that this tool will enable broad utilization of our data and will serve as a valuable repository for the broader EV community.

## Discussion

In our study, we integrated multi-omics investigation to systematically resolve the core protein and lipid componentry of circulating EVs in humans **(Figure 7)**. Our discovery includes a conserved set of 182 proteins and 52 lipids intrinsic to circulating EVs, and a panel of 29 proteins and 114 lipids that are NonEV features in plasma, which together serves as biological markers for EV research applicable to human samples. These extensive protein and lipid landscapes, which can be easily accessed with the Shiny web tool, will be instrumental to the EV community for clearer understanding of circulating EV biology. This includes the survey of circulating EV surfaceome, comprising of 151 EV protein features, that could be potentially exploited for antibody-based capture of circulating EVs based on their surface protein expression(s). In addition to this resource, we identified a minimal set of biomarkers - ADAM10 (protein), PS(36:1), and CE(18:0) (lipids) - with strong discriminatory power between EVs and NonEVs. This reduced marker set is compatible with targeted, scalable assays such as ELISA or targeted mass spectrometry, making it highly practical for clinical translation. Furthermore, the ranked feature list generated by our machine learning framework provides a valuable resource for the research community, enabling prioritization of alternative markers based on available reagents or disease-specific applications.

**Figure 7.**
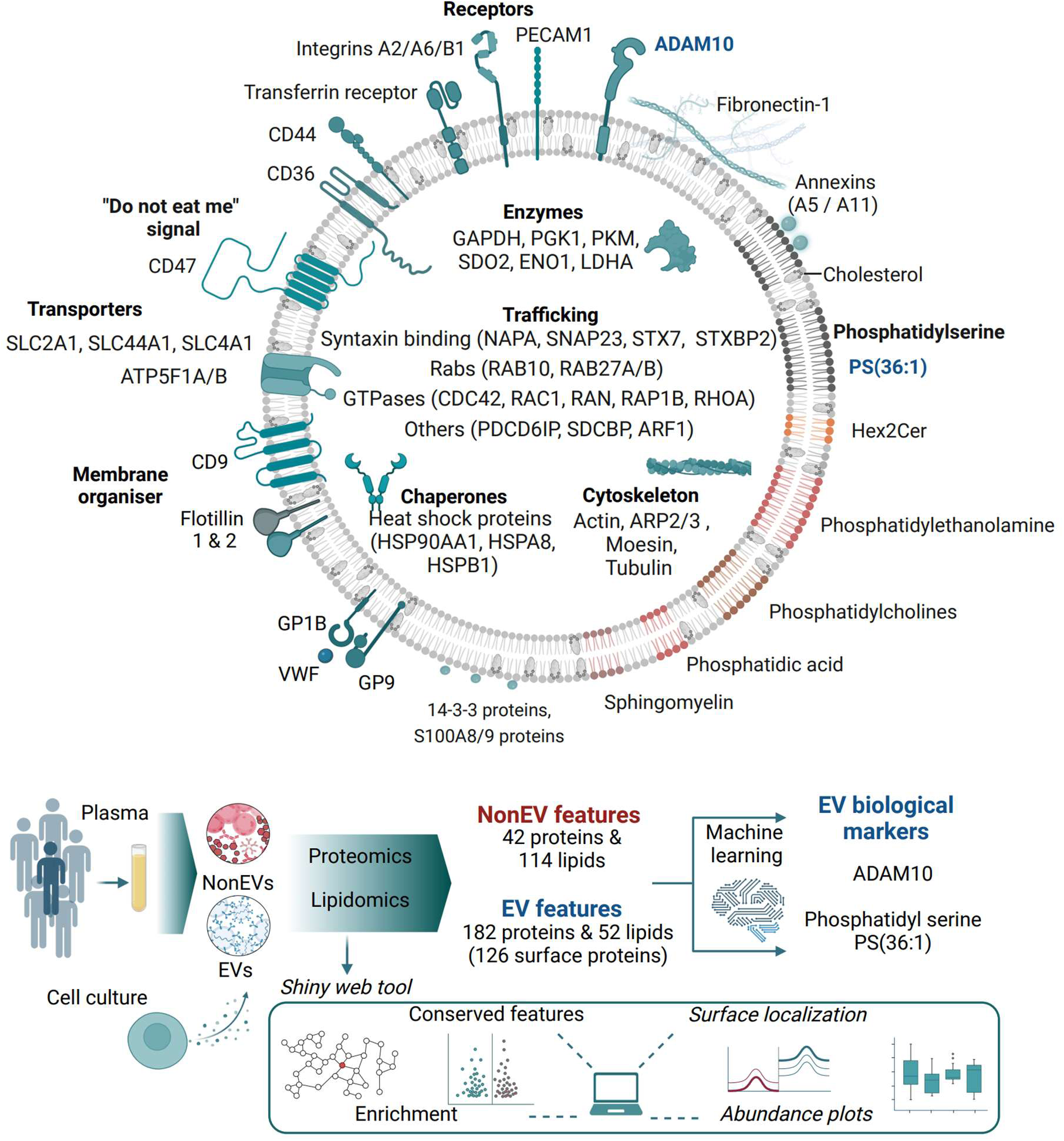
Hallmark molecular features of circulating EVs. **A.** Resolving the protein and lipid componentry intrinsic to circulating EVs. These features are also conserved in EVs released by cultured cells. We submit EV features, namely ADAM10 and phosphatidyl serine PS(36:1), as biological marker for confident EV identification and potentially purity assessment. The freely accessibly Shiny webtool will facilitate easy interrogation (conservation, abundance report, surface localization, and enrichments amongst others) of the multiomics data to encourage reuse.

The highly conserved protein ADAM10 on circulating EVs is also included in this EV surfaceome. The role of ADAM10 as conserved and robust EV marker is also supported by various studies that have reported ADAM10 expression in circulating EVs in human plasma, different biofluids, and EVs released by diverse tissue cell types using various biochemical analysis, different EV isolation strategies. Moreover, ADAM10 was also highly conserved in different circulating EV subpopulations (based on surface expression of tetraspanins, CD9, CD63, CD81^70^). The surface localization of ADAM10 is also corroborated in previous reports^88^. On the other hand, EV lipid features PS^15^ has already been developed for capturing circulating EVs using TIM4-based magnetic beads. The selective presence of ADAM10 (and PS lipids) in only a subset of EVs highlights the non-random nature of EV cargo packaging, supporting models of regulated biogenesis pathways such as ESCRT-dependent sorting^89^. Evidence that ADAM10 retains proteolytic activity in EVs^90^ suggests that these vesicles may serve as active mediators of extracellular remodeling and intercellular communication, expanding their functional relevance beyond passive biomarker carriers. The differential association of ADAM10 with specific EV subtypes, and its absence from some canonical small EV populations^87^, underscores the existence of biophysically and functionally distinct EV in circulation. Together, these findings support a growing consensus that EV heterogeneity is biologically regulated rather than stochastic, with specific protein profiles reflecting distinct functional roles in physiological and pathological contexts^87,89,90^.

Our data also provides “high-confidence” molecular leads for studying EV biology in multimodal organisms, particularly targeting EV biogenesis to curb EV-driven pathogenesis in complex organisms such as the mouse^83^ which remains largely unexplored. For instance, EV protein feature STEAP3 (also known as TSAP6) has been shown to regulate EV biogenesis in mice^91^. A holistic analysis of the core protein and lipid componentry can also provide insights into biological pathways co-involved in proteins and lipids; one such biology includes the potential involvement of lipid raft biology in human EVs. For instance, EV features such as flotillins^92^, SM and PS, along with cholesterol that are the most abundant lipids in EVs, are core components of lipid raft domains^84^. Assembly of flotillins in SM/ceremide^84^-rich microdomains induces curvature stress^93^ to regulate EV biogenesis and cargo selection^78,94^ in ESCRT^95^- and syntenin/syndecan-independent mechanisms^96^. Moreover, proteins that make the cortical actin (KEGG term enriched in EV core proteins) helps maintain and remodel these domains^84^. Accumulating evidence suggests that lipid rafts are nidus of EV biogenesis and are present in EVs as functional units^94^. Moreover, several bioactive cargo quantified in pEVs - such as TGFB1^66^, the RNA binding protein HNRNPK^62^, integrins^67^, SRC kinases^65^ - are closely linked to lipid raft assembly and trafficking^97–100^. Importantly, classical lipid raft–modulating proteins such as CAV1 and CAVIN1 - known to regulate membrane microdomain composition and influence the EV proteome^101,102^ - regulate miRNA cargo loading into EVs by modulating HNRNPK within the membrane raft^62^. Elevated levels of EV-associated HNRNPK have also been detected in body fluids from patients with metastatic prostate and colorectal cancers, underscoring its clinical relevance. Ubiquitous identification of these features in pEVs highlights similar EV biogenesis /cargo sorting mechanisms conserved for circulating EVs in humans.

A major caveat in our study is that it primarily focuses on bulk preparation of small EVs present in human plasma. Cells release heterogeneous EV populations^36,45,103,104^ that display functional diversity^105^. They can be categorized based on their origin^45,106,107^, density^45,50,104^, biochemical composition^31,50^ and size^47,50^. Since current EV isolation strategies cannot purify a specific subset of EVs to homogeneity, operational terms such as small EVs (30-150 nm) and large EVs (100-1000 nm) are encouraged where evidence for subcellular origin cannot be ascertained. In our study, small EVs potentially include both exosomes (endosomal-derived) and microvesicles (shed from plasma membrane) evident from our EV protein markers that include markers of both EV subtype (TSG101, ANXA2). Thus, our EV markers are applicable in a broad sense to small EVs; their specificity in exosomes *versus* microvesicles can potentially provide greater resolution in human EV biology and thus warrants future investigation. Moreover, EV heterogeneity is reported in circulating EVs too based on surface expression of tetraspanins, CD9, CD63, CD81^70^. Nonetheless, our EV protein features show high conservation in all three EV subpopulations^70^. While bulk EV study provides a broader understanding of the complex interactions and processes involving EVs within the body that specific EV subtypes might not fully capture, it is essential to study EV subpopulation-specific core features in humans, as this avenue holds great potential for advancing precise diagnostics and therapeutics. Nevertheless, bulk analyses can still offer insights into coordinated molecular patterns and biologically conserved pathways. For instance, enrichment patterns observed in our study (Extended DataL7A) reflects functionally associated protein modules enriched in the plasma EV pool. While these associations do not imply direct molecular interactions or spatial co-localization within individual EVs, they highlight coordinated expression patterns that are recurrent across EV subtypes and cell types. Importantly, we validated that many of these molecular features are enriched in EVs derived from multiple cell types - including cancer cells and primary fibroblasts - as well as across CD9-, CD63-, and CD81-positive plasma EVs, suggesting they are not artifacts of averaging but representative of fundamental EV-associated signatures. Our findings align with prior studies demonstrating that such molecules are actively enriched in EVs relative to their cells of origin and conserved across cell-derived EVs^30^. Thus, while single-vesicle resolution would offer even deeper insight, bulk EV analysis can still uncover biologically meaningful and source-representative features that contribute to systemic intercellular communication. In regards to source attribution, protein signatures associated with diverse cell types were represented in pEVs, including endothelial cells, fibroblasts, hepatocytes, cardiomyocytes, kidney cells, and hematopoietic cells (such as platelets), corroborating previous reports^15,54,55,108,109^. In healthy individuals, circulating EVs reportedly arise from hematopoietic cells (such as platelets, erythrocytes, and leukocytes) and endothelial cells^54,55^. While definitive source attribution of plasma EVs remains technically challenging due to the lack of universally exclusive surface markers, recent computational deconvolution algorithms offer promising solutions to estimate the relative contribution of different cell types^108,109^. Complementary proteomic studies have also identified tissue-specific EV-associated proteins, providing valuable leads for understanding EV origin and function in systemic circulation^15^. In parallel, high-sensitivity techniques such as single-vesicle flow cytometry and antibody-based surface profiling^110,111^ offer additional means to enrich and classify EVs based on cellular source, which may be used alongside our reference EV proteome dataset to further refine classification and improve translational application. Our dataset offers a high-confidence resource of tissue- and cell-enriched protein features in circulating EVs that can be harnessed to guide the development of affinity-based enrichment strategies, improve computational deconvolution models, and support source-specific EV biomarker discovery. In future studies, combining these molecular leads with advanced single-EV technologies and disease-specific profiling could yield deeper insights into tissue-specific EV biogenesis, intercellular communication, and their roles in health and disease. Importantly, despite disease-associated proteomic changes, the core set of 182 plasma EV protein features remained highly conserved across both CAC and non-CAC individuals in the EDCAD cohort, underscoring their robustness as molecular hallmarks of circulating EVs. This warrants future investigation on molecular stability across diverse disease states to establish the use of these core pEV features as a reliable foundation for human plasma EV studies in large population-based cohorts, enabling consistent characterization and cross-study comparability.

In summary, we identify core protein and lipid componentry of circulating EVs in humans. We submit these features as “hallmark molecular features” of circulating EVs that serve as valuable resource to the EV community and a tool to enhance the quality of human EV-research.

## Methods

### Generation of cell conditioned media

SW480 (CCL-288, ATCC), SW620 (CCL-227, ATCC) and LIM1863 cells^112^ (Ludwig Institute for Cancer Research, Melbourne) cells were cultured in RPMI-1640 (Life Technologies). MDA MB 231 (HTB-26, ATCC) was cultured in DMEM (Life Technologies). Primary human cell source include human neonatal foreskin fibroblast cell line (neoHFF) [sourced as gift from Pritinder Kaur, Monash University, Australia), human dermal fibroblasts, adult (Gibco/Thermo Fisher Sci. #C0135C) were cultured in Complete culture media included media supplemented with 5% (v/v) Fetal Bovine Serum (FBS, Life Technologies) and 1% (v/v) Penicillin Streptomycin (Pen/Strep, Life Technologies). Human atrial cardiac fibroblasts (Lonza, #CC-2903) and human ventricular cardiac fibroblasts (Lonza, #CC-2904) were cultured in FGM™-3 Cardiac Fibroblast Growth Medium-3 BulletKit™ (Lonza, #CC-3131 and #CC-4525). Human umbilical vein endothelial cells (HUVEC, Lonza, #CC-2519; HUVEC sourced as gift [Karlheinz Peter, BHDI, Australia], HUVEC-RFP, Angio-Proteomie #cAP-0001) were cultured in EGM™-2 Endothelial Cell Growth Medium-2 BulletKit™ (Lonza #CC-3156 and #CC-4176 supplements) prepared as detailed by supplier. All cells were cultured at 37 °C with 10% CO_2_ as described^45,68^. Cells were passaged with trypsin-EDTA (Gibco). Cells were cultured in CELLine AD-1000 Bioreactor classic flasks (Integra Biosciences) for conditioned media generation as described^68^.

### Plasma preparation

Human blood plasma samples were obtained from Australian Red Cross Lifeblood or EDCAD study. For Red Cross samples, ethical permits were obtained from the ethical committee of the Australian Red Cross Blood Service Human Research Ethics Committee, and La Trobe University Human Ethics Committee (HEC19485). Blood samples were collected via aseptic venipuncture into commercial EDTA-containing sampling containers at room temperature and centrifuged at 4,200 *g* for 10 min. The top plasma supernatant was carefully collected, and further centrifuged at 5,000 *g* (15 min) at room temperature. The supernatant was immediately isolated and stored as 1 ml aliquots at -80°C until further use. For EDCAD samples, ethics permit was approved through Human Research Ethics Committee (HREC) at Baker Heart and Diabetes Institute and by the Alfred Hospital Ethics Committee (EDCAD-PMS, #492/20). Blood was collected via venipuncture into a sterile EDTA tube and centrifuged at 2,700 g at room temperature for 13 min. A small volume (1.6 ml) of the supernatant (plasma) was transferred into a clean sterile tube and Butylated Hydroxytoluene solution (BHT) added to a final concentration of 100 µM. The Plasma/BHT sample was mixed and equally transferred into 3 x 0.75 ml Fluid X cryotubes. Plasma/BHT samples were snap frozen on dry ice prior to being transferred and stored at -80°C until use). For AusDiab samples, ethics permit was approved through HREC at Baker Heart and Diabetes Institute and by the Alfred Hospital Ethics Committee (#39/11). Blood was collected into commercial fluoride/oxalate tubes and centrifuged at 2,500 rpm for 10 min at room temperature. Plasma was isolated and snap frozen on dry ice prior to being transferred and stored at -80°C until use. Clinical parameters for EDCAD plasma samples are provided in **Supplementary Table 41**.

For all plasma collections, only ∼80% of top plasma supernatant was collected to avoid disturbing the buffy coat. Samples with potential hemolysis, cellular contamination and high fat/lipid content were not used for this study.

### Ultracentrifugation of plasma or conditioned media

To generate crude 100K plasma pellet, plasma samples were thawed on ice and subjected to ultracentrifugation at 100,000 *g* for 1 hr or 18 hr at 4 °C (TLA-55 rotor; Optima MAX-MP Tabletop Ultracentrifuge, Beckman Coulter). Pellets were washed once in 1.0 ml PBS, resuspended in 100 µl PBS and stored at -80°C until further use.

Human fibroblasts and endothelial cells (80% confluent in T75 culture flasks) were cultured in DMEM supplemented with 0.5%v/v insulin transferrin selenium (Invitrogen) and 1% Pen/Strep) to generate conditioned media. To isolate EVs, we subjected conditioned media to differential centrifugation (500 *g* for 5 min and 2,000 *g* for 10 min at 4 °C), followed by concentration using Amicon Ultra Centrifugal Filter, 100 kDa MWCO (low speed centrifugation at 2,000 *g* at 4 °C). The filter was washed with 5 time with 1 ml ice-cold PBS. The concentrated retentate was retrieved and further centrifuged at 10,000 *g* (30 min at 4 °C) to pellet crude large EVs (10K EVs), while the resultant supernatant underwent ultracentrifugation at 100,000 *g* (1 h at 4 °C) to pellet crude small EVs (100K EVs). The isolated EVs were washed in PBS (1 ml, 10,000 *g* for large EVs and 100,000 *g* for small EVs) before resuspension in 50 µl PBS and stored at -80°C until further use (subsequent proteomic and lipidomic analyses).

### Direct density-gradient separation

Plasma (500 µl) was directly overlaid on top of discontinuous gradient of OptiPrep^TM^ (40% (3 ml), 20% (3 ml), 10% (3 ml) and 5% (2.5 ml) (diluent: PBS solution)) and ultracentrifuged at 100,000 *g* for 18 hr (4 °C, 41 Ti rotor; Optima XPN Ultracentrifuge). Twelve 1 ml fractions were carefully collected top-bottom, diluted in PBS (2 ml) and ultracentrifuged at 100,000 *g* (1 h, 4°C, TLA-55 rotor; Optima MAX-MP Tabletop Ultracentrifuge). Fraction densities were determined as previously described^45^. Pellets were suspended in 100 µl PBS and stored at -80 °C until further use. For bottom-up DGS, 500 µL of plasma was carefully loaded at the bottom of a pre-formed discontinuous DGS (OptiPrep^TM^ (40% (3 ml), 20% (3 ml), 10% (3 ml) and 5% (2.5 ml) (diluent: PBS solution)) using a long syringe to avoid disturbing the gradient. Ultracentrifugation was performed at 100,000 *g* for 18 h (4°C), as in the top-loaded DGS method. After centrifugation, 12 equal fractions (1 ml each) were collected, diluted with 2 ml PBS, and subjected to another round of ultracentrifugation at 100,000 *g* for 1 h (4°C). The resulting pellets were resuspended in 50 µl PBS and analysed by Western blotting.

For isolation of cell culture-derived EVs, conditioned media was centrifuged at 500 *g* (5 min, 4 °C), 2,000 *g* (10 min, 4 °C), and 10,000 *g* (30 min, 4 °C, SW28 rotor; Optima XPN Ultracentrifuge, Beckman Coulter) to pellet large EVs^68^. The supernatant (1 ml) was subjected to direct DGS for EV isolation as described for plasma above. EV pellets were reconstituted in 100 µl PBS and stored at -80 °C until further use.

### Immunoblotting

Samples were subjected to protein quantification (microBCA™ Protein Assay Kit (23235, Thermo Fisher Scientific)), Western blotting (iBlot 2 Dry Blotting System, Thermo Fisher Scientific) was performed as described^45,68^. Rabbit antibodies raised against, Albumin (ab207327, Abcam), AGO2 (ab186733, Abcam) and Fibronectin (ab2413, Abcam) were used. Mouse antibodies CD63 (556019, BD Pharmingen™), CD81 (555675, BD Pharmingen™), APOB100 (3715-3-250, Mabtech), APOA1 (3710-3-1000, Mabtech) and ALIX (2171S, CST) were used. Secondary antibodies used were IRDye 800 goat anti-mouse IgG or IRDye 700 goat anti-rabbit IgG (1:15000, LI-COR Biosciences).

### Biophysical characterization of EVs

Cryo-electron microscopy (Tecnai G2 F30) on samples (1 µg) was performed as described^47^. Briefly, Plasma 100K, D-DGS fractions 1-3 (pooled) and Plasma EVs (DGS fractions 6-8, pooled) (∼1 µg) were loaded onto to glow-discharged C-flat holey carbon grids (ProSciTech Pty Ltd) that, after blotting away excess liquid, were plunge-frozen in liquid ethane and subsequently mounted in a Gatan cryoholder (Gatan, Inc.) in liquid nitrogen. Images were then acquired at 300 kV using a Tecnai G2 F30 (FEI) in low-dose mode. EV samples (∼1 µg; 1:1000 dilution) were prepared in PBS (#14190-144, Thermo Fisher Scientific)^113^ and particle size distribution and zeta potential (surface charge) determined by nanoparticle tracking analysis (ZetaView, Particle Metrix, PMX-120; 405 nm laser diode) according to manufacturers’ instructions.

### Surface biotin-labelling of EVs and proteomic sample preparation

EV surface proteins were captured using biotinylated using Pierce™ Cell Surface Biotinylation and Isolation Kit (A44390, Thermo Fisher Scientific), digested with trypsin (Promega, V5111), and peptides analyzed using Q-Exactive HF-X as described^68^.

### Label-free proteomics sample preparation

Label-free proteomics sample preparation (∼2-10 µg in 20-50 µl) was performed as described^68^, using single-pot solid-phase-enhanced sample preparation (SP3) workflow^114^. Briefly, samples were solubilized (1% (v/v) sodium dodecyl sulphate (SDS), 50 mM HEPES pH 8.0), incubated at 95 °C for 5 min and cooled. Samples were reduced with 10 mM dithiothreitol (DTT) for 45 min at 25 °C followed by alkylation with 20 mM iodoacetamide for 30 min at 25 °C in the dark. The reaction was quenched to a final concentration of 20 mM DTT. Magnetic beads were prepared by mixing SpeedBeads™ magnetic carboxylate modified particles (65152105050250, 45152105050250, Cytiva) at 1:1 (v:v) ratio and washing twice with 200 µl MS-water. Magnetic beads were reconstituted to a final concentration of 100 µg/µl. Magnetic beads were added to the samples at 10:1 beads-to-protein ratio, and 100% ethanol (EA043, ChemSupply) added for a final concentration of 50% ethanol (v/v). The sample bead mix were incubated at room temperature with constant shaking (1000 rpm). Protein-bound magnetic beads were washed three times with 200 µl of 80% ethanol and reconstituted in 50 mM TEAB and digested with Trypsin and Lys-C (1:50 and 1:100 enzyme-to-protein ratio, respectively) for 16 h at 37 °C with constant shaking (1000 rpm). The peptide mixture was acidified to a final concentration of 2% formic acid (pH <2) and centrifuged at 20,000 *g* for 1 min. The peptide digests were frozen at -80°C and dried by vacuum centrifugation (Savant SPD121P, Thermo Fisher Scientific), reconstituted in 0.07% trifluoroacetic acid, and quantified by Fluorometric Peptide Assay (23290, Thermo Fisher Scientific) as per manufacturer’s instruction.

### TMT-based proteomics sample preparation

For TMT-based labelling was performed as described^115,116^, using 10-plex TMT according to manufacturer’s instructions with minor modifications (Thermo Fisher Scientific, 90406, lot UG287488/278919). The 10-plex experiment comprised 9 different chemical tags for EV peptide labelling, with the tenth tag used for generating reference channel made by a pooled peptide digest from all samples. A list of the sample labelling strategy is available in MassIVE proteomeXchange (MSV000094307). Peptide samples (15.5 µg) were labelled with TMT10plex™ Isobaric Label (90406, Thermo Fisher Scientific) at 4:1 label-to-peptide ratio for 2 hr at room temperature and quenched with 0.5% (v/v) hydroxylamine for 30 min at room temperature. Labelled peptide samples were acidified with 4% (v/v) formic acid (FA) and pooled into a new microtube. Pooled samples were desalted using Sep-Pak tC18 96-well µElution (186002318, Waters) and elution lyophilized by vacuum-based speedVac. Peptides were then fractionated into 20 fractions (with increasing concentration of ACN from 2-50%) using high pH reversed-phase chromatography using in-house SPE StageTips (SDB-RPS, Empore) as described^117^. Peptide eluates were lyophilized by vacuum-based SpeedVac, reconstituted in 0.07% TFA and quantified using Colormetric peptide assay. Peptide fractions with low yields were pooled.

### Nano liquid chromatography–tandem mass spectrometry

Peptides (label-free and TMT-labelled) were analyzed on a Dionex UltiMate NCS-3500RS nanoUHPLC coupled to a Q-Exactive HF-X hybrid quadrupole-Orbitrap mass spectrometer equipped with nanospray ion source operating in positive mode as described.^68,115,118,119^

For label-free proteomics, peptides were loaded (Acclaim PepMap100 C18 3 µm beads with 100 Å pore-size, Thermo Fisher Scientific) and separated (1.9-µm particle size C18, 0.075 × 250 mm, Nikkyo Technos Co. Ltd) with a gradient of 2–28% acetonitrile containing 0.1% formic acid over 49 or 95 mins at 300 nl min^-1^ at 55°C (butterfly portfolio heater, Phoenix S&T). A list of the run details is available in MassIVE proteomeXchange (MSV000094307).

For data-dependent acquisition (DDA), MS1 scan was acquired from 350–1,650 *m*/*z* (60,000 resolution, 3 × 10^6^ automatic gain control (AGC), 128 msec injection time) followed by MS/MS data-dependent acquisition (top 25) with collision-induced dissociation and detection in the ion trap (30,000 resolution, 1 ×10^5^ AGC target, 27 msec injection time, 28% normalized collision energy, 1.3 *m*/*z* quadrupole isolation width). Unassigned, 1, 6-8 precursor ions charge states were rejected and peptide match disabled. Selected sequenced ions were dynamically excluded for 30 s. Data was acquired using Xcalibur software v4.0 (Thermo Fisher Scientific).

For DIA^120^, full scan MS were performed in the m/z range of 350 to 1100 m/z with a 60,000 resolution, using an automatic gain control (AGC) of 3 x 10^6^, maximum injection time of 50 ms and 1 microscan. MS2 was set to 15,000 resolution, 1e6 AGC target and the first fixed mass set to 120 m/z. Default charge state set to 2 and recorded in centroid mode. Total scan windows (38 windows; 49 min gradient, 63 windows, 95 min gradient) with staggered isolation window from 350 to 1100 m/z were applied with 28% normalized collision energy as described^121^.

TMT labelled peptides were separated (1.9-µm particle size C18, 0.075 × 250 mm, Nikkyo Technos Co. Ltd) with a gradient of 4-28% ACN containing 0.1% FA over 110 min at 300 nL/min at 55 °C (butterfly portfolio heater, Phoenix S&T). The MS1 scan was acquired from 300 to 1650 m/z (60,000 resolution, 3e6 AGC, 128 ms injection time) followed by MS/MS data-dependent acquisition of the top 15 ions with HCD (30,000 resolution, 1e5 AGC, 60 ms injection time, 33 NCE, 0.8 m/z isolation width). Unassigned, 1, 6-8 precursor ions charge states were rejected and peptide match disabled. Selected sequenced ions were dynamically excluded for 30 s.

MS-based proteomics data (including sample/label annotation) is deposited to the ProteomeXchange Consortium via the MassIVE partner repository and available via MassIVE with identifier (MSV000094307).

### Proteomic data processing

For DDA and TMT proteomics, peptide identification and quantification were performed using MaxQuant (v1.6.6.0-v1.6.14) with its built-in search engine Andromeda as described^68^ against *homo sapiens* (UP000005640) including contaminants database. For LFQ-based analyses (DDA), cysteine carbamidomethylation was selected as a fixed modification and N-terminal acetylation and methionine oxidations as variable modifications. Data was processed using trypsin/P as the proteolytic enzymes with up to 2 missed cleavage sites allowed. Further processing using match between runs (MBR) and label free quantification (LFQ) algorithm were employed. Peptides were identified with an initial precursor mass deviation of up to 7 ppm and a fragment mass deviation of 20 ppm with a minimum length of 7 amino acids. False discovery rate (FDR) was 0.01 for both the protein and peptide by searching against a reverse database. For TMT-based analyses, reporter ion MS2 (TMT10plex) settings were employed. For biotin surface proteome analysis additional Thioacyl (DSP) (C(3)H(4)OS)) was employed^47^.

For DIA proteomics, data processing was performed using DIA-NN^122^ (v1.8). Spectral libraries were predicted using the neural network deep learning algorithm employed in DIA-NN with Trypsin/P, allowing up to 1 missed cleavage. The precursor change range was set to 1-4, and the m/z precursor range was set to 300-1800 for peptides consisting of 7-30 amino acids with N-term methionine excision and cysteine carbamidomethylation enabled as a fixed modification with 0 maximum number of variable modifications. The mass spectra were analyzed using default settings with a false discovery rate (FDR) of 1% for precursor identifications and match between runs (MBR) enabled for replicates. The resulting output files contaminants and reverse identifications were removed.

### Lipid extraction

Lipids were extracted from samples using a modified single-phase butanol/methanol extraction method as described^123^. In brief, the samples (2-5 μg protein from lyophilized sample reconstituted in 10ul of water) were mixed with 100 μl of 1:1 butanol:methanol containing the internal standards, the samples were vortexed, sonicated on a sonicator bath (1 h) and centrifuged (13,000L*g*, 10 min). The supernatants were transferred into glass vials and stored at −80 °C. Prior to mass spectrometry analysis, samples were thawed for 1Lhr at room temperature, vortexed and sonicated on the sonicator bath for 15 min and left to sit at 20L°C for 2 h prior to analysis.

### Liquid chromatography mass spectrometry (discovery)

Analysis of lipid extracts was performed on an Agilent 6495C QQQ mass spectrometer with an Agilent 1290 series HPLC system. Mass spectrometry settings and transitions for each lipid class are provided (**Supplementary Table 40**) adapted from previous methodology^124^. Mass spectrometer running conditions were gas temperature 150 °C, gas flow rate 17Ll/min, nebulizer gas pressure 20Lpsi, sheath gas temperature 200 °C, capillary voltage 3500LV and sheath gas flow 10Ll/min. Isolation widths for Q1 and Q3 were set to “unit” resolution (0.7Lamu). The Agilent 1290 Infinity II HPLC conditions were as follows: The composition of running solvents A and B comprised 50:30:20 and 1:9:90 water, acetonitrile and isopropanol, respectively. Solvent A was buffered with 10mM ammonium formate with medronic acid, while Solvent B was buffered with 10mM ammonium formate. A ZORBAX eclipse plus C18 (2.1L×L100Lmm 1.8Lµm, Agilent) column was used. Solvents were run at a flow rate of 0.4 ml/min with the column compartment temperature set to 45 °C.

The solvent gradient was as follows: starting at 15% B and increasing to 50% B over 2.5Lmin (while diverting to waste for the first min), then to 57% B over 0.1 min, to 70% B over 9.4 min, to 80% B over 0.5 min, to 90% B over 3.5 min and finally to 100% over 0.1Lmin. The solvent was then held at 100% B for 0.9 min and returned to 15% B over 0.1Lmin and held for 2.9 min (a combined total of 20 min run time).

### Liquid chromatography mass spectrometry (validation) for AusDiab and fibroblasts/endothelia cell EVs

Analysis of lipid extracts was performed on an Agilent 6495C QQQ mass spectrometer with an Agilent 1290 series HPLC system similar to the discovery analysis. Optimized conditions to increase sensitivity was used. Mass spectrometer running conditions were gas temperature 200L°C, gas flow rate 17Ll/min, nebulizer gas pressure 20Lpsi, sheath gas temperature 280L°C, capillary voltage 3500 V and sheath gas flow 10Ll/min. Isolation widths for Q1 and Q3 were set to “unit” resolution (0.7Lamu). Solvent composition was identical to the discovery run. The solvent gradient was as follows: starting at 15% B and increasing to 50% B over 2.5 min (while diverting to waste for the first min), then to 57% B over 0.1 min, to 70% B over 6.4 min, to 93% B over 0.1 min, to 96% B over 1.9 min and finally to 100% over 0.1 min. The solvent was then held at 100% B for 0.9 min and returned to 15% B over 0.1Lmin and held for 3.8 min (a combined total of 16 min run time per sample). Additional characterisation was carried out on PC, PC(O) and PC(P) isobaric and isomeric species to highlight the separation of the chromatography (**Supplementary** Figure 20).

### Lipid data integration and statistical analysis

Raw MS data was analyzed using MassHunter Quant 10.0 (Agilent Technologies). Peak area of the lipid species was normalized to their respective internal standards to generate relative concentration data per sample. For lipids that appear to have isomeric separation on the chromatography, they are designated with the (a) and (b) annotations to highlight different elution orders. Lipid class totals were generated by summing the individual species within each class. MS-based lipidomic sample/label annotation is deposited to the ProteomeXchange Consortium via the MassIVE partner repository and available via MassIVE with identifier (MSV000094307).

### Structural characterisation of lipid isomers and isobars

As PS 36:1 represented a strong signature for EVs, we determined the structure of this PS species in isolated EV’s, we re-ran pooled samples under negative ionisation mode under the same gradient conditions screening for product ions corresponding to the serine head group (788.5 m/z -> 701.5 m/z) and the fatty acyl tails. The final composition was annotated to be PS 18:0_18:1 with signals observed with 283.3 and 281.3 product ions (**Supplementary** Figure 19). The method run for these samples monitors CE’s in positive ionisation mode as an ammonium adduct monitoring for the cholesterol ion with additional water loss (CE 18:0 [M+NH4]+ 670.6 / 369.3). We have a matching internal standard (CE 18:0 d6) to provide quantitation for this lipid species. Retention time, mass and fragmentation confirms the current annotation.

### Nanoflow analysis of plasma EVs

Plasma EVs (∼5 µg) were subjected to fixation and permeabilization using eBioscience™ Foxp3 / Transcription Factor Staining Buffer Set (Invitrogen™, 00-5523-00). Briefly, pEVs (in 50 µl PBS) was incubated with 500 µl fixation and permeabilization buffer on ice for 30 mins. Samples were ultracentrifugation at 100,000 g (1 h at 4 °C) and pellets resuspended in 100 µl wash buffer. Samples were stained with 5 µl APC Annexin V reagent (BioLegend) and either 2 µg of Anti-ADAM 10 Antibody (Sigma-Aldrich, AB19026) or Rabbit IgG Isotype Control (Invitrogen, 10500C). Samples were incubated at room temperature (gentle end-over mixing) for 1h. Samples were topped with 900 µl wash buffer and washed twice (ultracentrifuged at 100,000 g (1 h at 4 °C)) to remove any remaining antibodies and potential antibody aggregates. The pellets resuspended in 100 µl wash buffer containing 0.5 µl of Goat Anti-Rabbit Alexa Fluor 568 Dye secondary antibody (ThermoFisher Scientific) incubated for 30 mins at room temperature (gentle end-over mixing) in dark. Samples were topped with 900 µl wash buffer and washed twice (ultracentrifuged at 100,000 g (1 h at 4 °C)). Pellets resuspended in 100 µl of PBS (0.5% bovine serum albumin) filtered using 0.22 µm filter. Samples were immediately analysed using Cytek Aurora flow cytometer and SpectroFlo^®^ software, as previously described^86^. Briefly, controls included isotype control antibodies were used to test for unspecific binding, in addition to unstained EV suspension, and buffer-only controls. For purity assessment, Sodium dodecyl sulfate detergent (SDS) was added to labelled EVs at a final concentration of 0.5%, followed by vigorous vertexing for ≈30 s before acquisition. High purity would be indicated by a strong reduction in vesicle concentration and fluorescence after the treatment. The instrument setLup was consistent across all experiments and followed recommendations from the manufacturer. Prior to the acquisition, the flow cell was washed with Contrad to minimize machineLassociated noise. Instrument gating calibration was performed using 90 nm (#64009-15) 125 nm (#64011-15), 150 nm (#64012-15), 200 nm (#64013-15) and equal mix (90-200 nm) beads (Nanobead NIST Traceable Particle Size Standards). The threshold for side scatter was set to 430, and the gain of side scatter (SSC) were set to 10. YG3-A channel used for detecting Alex Flour 568 signal, R1-A channel used for detecting APC signal and 10 000 events were recorded for all samples with the slowest flow rate to minimize the swarming effect. Statistical analysis of flow cytometry values was performed using GraphPad (v 9.1.0) using multiple paired t-tests.

### Isolation of extracellular vesicles using affinity to TIM4

To capture PS-positive EVs from plasma, human TIM4-Fc protein (FUJIFILM Wako Pure Chemical Corporation) was biotinylated with EZ-Link™ Maleimide-PEG11-Biotin (Thermo Fisher Scientific) and conjugated to Dynabeads™ MyOne™ Streptavidin C1 magnetic beads (65001, Thermo Fisher Scientific) as described^125^. Briefly, plasma (200 µL), diluted with 300 µl of PBS, was supplemented with heparin sodium (4U, Thermo Fisher Scientific) and CaCl_2_ (final concentration of 2 mM) and incubated with TIM4 affinity beads for 16 hr at 4°C with gentle rotation. Beads were collected and washed 5x with 1 mL Tween-TBS (0.05% Tween-20, 2 ml CaCl_2_). EVs were eluted in 50 µl PBS containing 1 mM EDTA and subjected to proteomic analyses using Q-Exactive HF-X as described above.

### Data processing and statistical analysis

Software tools used for this study are freely available as open-source R packages (https://www.r-project.org). No newly generated software or custom code were used in this current study and hence the codes have not been deposited in a public repository but is available from the corresponding author(s) upon request.

For key analyses, proteome and lipidome data sets were analyzed using R package Differential Enrichment analysis of Proteomics data (DEP)^126^. Proteins identified in at least 70% of one biological group were selected for downstream analysis. Using DEP, the data was background corrected and normalized by variance stabilizing transformation (vsn, which also log2-transforms the data), followed by imputation of missing values, whereby missing at random data were ‘knn’ imputed and missing not at random data were ‘MinProb’ imputed. Protein-wise linear models combined with empirical Bayes statistics were used for the differential enrichment analysis, whereby the raw p-values were adjusted to correct for multiple testing using Benjamini-Hochberg method. Differentially abundant proteins or lipids were clustered by k-means clustering using DEP package. The PCA plot, Pearson Correlation matrix, volcano plots, log2 centred bar plots and overlap bar plots were also generated using DEP. Heatmaps were generated using ComplexHeatmap package^127^. Box plots and scatter plots were generated using RStudio package ggplot2. Cytoscape^128^ was used to generate Ontology map^129^ (plugin v3.7.1). Bioconductor package clusterProfiler 4.0^130^ was used to perform Ontology or KEGG pathway enrichment analysis, or gene set enrichment (KEGG) analysis, with default parameters used to identify significantly enriched gene sets. The pathway-based data integration and visualization was constructed using R package pathview^131^. For identification of lipid-associated terms enriched in lipidomes, LION web-based ontology enrichment tool^132^ was used.

For conserved protein identification analysis, proteins were classified as detected or not detected across samples (protein abundance was not considered). In surface biotin-labelling of EVs, proteins detected in at least 6 out of 9 experiments were considered as pEV surface proteins (protein abundance was not considered). For annotating surface proteins into different categories, SURFY ^59^-based categorical annotation of cell surface proteins were employed. To identify pEV lipid features, we performed K-means clustering of differentially abundant lipids between pEVs *versus* NonEVs and in vitro EVs *versus* cells using DEP package.

We employed caretEnsemble R package to assess the ability of these protein features (182 pEV protein features and 42 NonEV protein features, using Naïve Bayes algorithm), surface protein features (comprising of 151 surface proteins associating with pEV surface and 32 surface proteins associating with NonEVs, using Naïve Bayes algorithm) or lipid features (114 pEV lipid features and 52 NonEV lipid features, using Neural Network algorithm (’nnet’)) to distinguish between distinguish between pEV and NonEV particles. The pEV and NonEV proteome/lipidome datasets were evenly partitioned based on sample type into training set (70% of the samples) and a validation set (remaining 30% of samples). For protein features based machine learning, proteomes of pEVs (n=16) and pDGS.LD (NonEV particles, n=16) from as 16 plasma samples of the EDCAD cohort served as independent test set. For lipid features based machine learning, lipidomes of DGS plasma fractions served as independent test set. A bootstrapping resampling method was employed with 25 resampling iterations to train the models. The performance of the models were evaluated using class probabilities and the two-class summary function. The preprocessing steps of centering and scaling were applied to the predictors before training the models. A confusion matrix that compare predicted classes to actual classes based on machine learning algorithm used were generated to visualize the predictive performance of the model. For identification of ADAM10 protein or PS(36:1) and CE(18:0) lipids as biomarker candidates, Recursive Feature Elimination (RFE) provided by the caret R package for feature selection using default options was used.

To address the question of source attribution in a data-driven manner, we manually curated tissue- and cell-type-specific protein signatures from three independent sources: (1) The Human Protein Atlas (HPA), which provides tissue- and organ-specific proteins based on immunohistochemistry **(Extended Data 2A-B)**; (2) HPA nTPM dataset, an RNA-derived protein expression database across tissues and cells **(Extended Data 2C)**; and (3) Organ-specific protein signatures detected in human plasma, identified via aptamer-based detection^57^ (**Extended Data 2D**). To systematically assess the presence of bioactive molecules in pEVs, we manually curated and integrated multiple independent datasets that catalogue biologically significant protein classes. Specifically, we leveraged established resources for human transcription factors^61^ (Human TFs Database), RNA-binding proteins^60^ (RBPDB Database), kinases^58^ (KinHub Database), and cell-surface receptors, transporters, and signalling proteins^59^. In addition, we incorporated Gene Ontology terms related to cytokine activity (GO:0005125), chemokine activity (GO:0008009), growth factor activity (GO:0008083), and signal transduction (GO:0007165) to ensure comprehensive coverage of bioactive EV-associated molecules.

The Shiny web application (https://evmap.shinyapps.io/evmap/) powered by R and hosted on shinyapps.io, was created using the R packages shiny, gplots, and ComplexHeatmap, and offers feature selections and visualizations for EV protein and lipid feature conservations in circulating EVs.

### Statistics and reproducibility

At least three independent biological replicates were used to generate the data. The raw p-values were adjusted to correct for multiple testing using Benjamini-Hochberg method and reported in the corresponding figure legends and supplementary tables.

### Data Availability

The data supporting the findings of this study are available within the manuscript, as supplementary information files or as source data files. The raw MS files and the search/identification files obtained using MaxQuant/DIA-NN have been deposited to the ProteomeXchange Consortium via the MassIVE partner repository and available via MassIVE with identifier (MSV000094307).

*Reviewer access to data repository can be made through MassIVE; MSV000094307 and reviewer portal:* ftp://MSV000094307@massive.ucsd.edu

## Acknowledgements

D.W.G laboratory is supported by research funds from National Health and Medical Research Council (NHMRC, MRF2015523, APP1141946), National Heart Foundation (NHF, 105072), Helen Amelia Hains Fellowship (D.W.G) and Department of Defense (PR230065). The Baker Heart & Diabetes Institute acknowledges support by the Victorian State Government Operational Infrastructure funding. We thank Eric Hanssen and the Bio21 Molecular Science and Biotechnology Institute for assisting cryo-electron microscopy (University of Melbourne). K.H is supported by NHMRC Emerging Leadership Fellowship (GNT1197190). Q.H.P, H.F and B.C are supported through an Australian RFP Scholarship, in addition to Bright Sparks Foundation. All figures were made in BioRender.com.

## Author Contribution

A.R. and D.W.G conceptualized the idea. A.R. designed the experiments and wrote the manuscript. A.R., Q.H.P., H.F., B.C., J.C., T.D., C.D., and D.W.G. performed experiments. A.R. performed bioinformatics analysis. A.R., K.H., and D.W.G., analyzed the results. C.D., J.E.S., and T.H.M., provided resources. A.R., K.H, B.C., J.E.S., T.H.M., P.M., and D.W.G. reviewed manuscript for submission.

## Supplementary Figure Legends

**Supplementary** Figure 1**. Characterization of EVs from human plasma**. **A**. Ultracentrifugation of human plasma. The protein yields of plasma and isolated EV pellets are indicated. Western blot analysis of the crude EV pellets for indicated proteins. **B.** Cryo EM image-based size distribution of plasma EVs and previously reported sizes for small (s-EV) and large EVs (L-EVs)^68^. **C**. Protein yield (µg) of pEVs per ml of plasma. **D.** Nanoparticle tracking analysis based total particle counts in DGS fractions (1-3) and DGS fractions (6-8) fractions/pEVs per ml of plasma (n=7). **E**. Comparative summary of biophysical characterization of pEVs and NonEVs. **F.** Bottom-up density gradient separation (DGS) of human plasma (0.5 ml). Western blot analysis of twelve DGS fractions (vol:vol matched) of human plasma with antibodies against indicated proteins.

**Supplementary** Figure 2**. Construction and characterization of circulating EV proteome landscape**. **A**. Workflow depicting sample details and MS-acquisition modes for constructing proteome landscape of plasma EVs (pEVs). Different experimental groups are indicated as Set, totalling six sets. **B**. Classification of different proteomes (from sets 1-5) into pEVs or nonEV groups. Bar plot depicts number of proteins identified within each proteome. Box plot represents normalised protein abundance (log10 intensity). **C.** Venn diagram of proteins identified indicated proteomes. **D.** Protein ranked by their abundance within indicated proteomes in Set 1. Proteins typical reported as EVs proteins are highlighted. **E.** Correlation of proteins in plasma, pDGS.LD and pEVs proteomes (median protein intensities of replicates) to the published concentration of the same proteins^49^. The black lines are linear regression models (the grey shaded regions represent 95% confidence interval). **F.** Heatmap of Pearson correlation matrix of protein abundance between proteomes.

**Supplementary** Figure 3**. Classical hallmark features of EVs preserved in circulating EV proteome**. **A.** Heatmap of differentially abundant proteins (p<0.05, log2 FC >1.5) with k-means clustering. **B.** KEGG pathways and Gene Ontology pathways enriched in each cluster (p<0.05) **C.** Leading edge analysis indicated for selected GSEA-KEGG pathways enriched in pEVs vs NonEVs. **D.** Bar plots showing relative abundance (log2 centred MS-based intensities) of differentially abundant EV and abundant plasma proteins.

**Supplementary** Figure 4**. Quantification of platelet EV proteins in circulating EV proteome**. **A.** Detection of platelet proteins (CLEC1B, PF4, PPBP) in platelet EVs in our previous report^53^. B. Detection of platelet EV proteins (CLEC1B, PF4, PPBP) in in vitro EV, NonEV or pEV proteomes. C. Detection of platelet EV proteins (CLEC1B, PF4, PPBP) in non-transformed cell-derived EVs. D. Detection of platelet EV proteins (CLEC1B, PF4, PPBP) in pEVs and NonEV proteomes from AusDiab Validation set.

**Supplementary** Figure 5. **Mapping EV markers in circulating EV proteome**. **A.** Pair-wise comparison of differentially abundant proteins (p<0.05, FC > 1.5) depicted as a volcano plot where 22 core proteins established for in vitro EVs^30^ and ninety-four EV marker proteins recommended by MISEV^36^ guidelines are depicted. **B**. Heatmap showing quantification of current EV marker proteins in pEVs. **C**. Occurrence analysis for current EV marker proteins in 38 pEV proteomes. **D**. Heatmap of selected EV proteins depicting their identification and abundance in 38 pEV proteomes.

**Supplementary** Figure 6. **Surfaceome of circulating EVs in human plasma. A.** Workflow for capture and identification of EV surface proteins. **B.** Categorization of pEV surface proteins and their relative abundance in pEVs vs pDGS.LD proteomes. **C.** Surface-accessible Category 1-3 proteins of pEVs categorised based on their molecular/functional annotation. C/R/T represents cluster of differentiation (CDs), receptors and transporters.

**Supplementary** Figure 7. **Conservation of circulating EV protein features in cell culture-derived EVs published proteomes. A.** Relative abundance of EV protein features in sEVs vs their parental cell proteome as reported^30^, whereby EVs were isolated from culture media using ultracentrifugation (UC). **B.** Relative abundance of EV protein features in sEVs vs their parental cell proteome as reported^30^, where EVs were isolated using density gradient separation (DG), size-exclusion chromatography (SEC), and UC.

**Supplementary** Figure 8**. Proteome analysis of pEVs, NonEVs and neat plasma from EDCAD set.** Iindividuals with either positive CAC score (CAC group) or zero CAC score (Healthy group) are indicated. **A**. Proteins quantified in pEVs, NonEVs and neat plasma from individuals with either positive CAC score (CAC group) or zero CAC score (Healthy group). **B.** Box plot depicting normalised intensities (Z-scored) of indicated proteins. **C.** Heatmap of Pearson correlation of quantified proteins. **D.** Principal component analysis of quantified proteins.

**Supplementary** Figure 9**. Lipidome profiling of circulating EVs. A.** Lipid abundance (MS-based intensity) for in vitro EVs, pEVs, p100K, pDGS.LD, and cells (WCL). **B.** Heatmap of Pearson correlation matrix of lipid abundance between datasets.

**Supplementary** Figure 10**. Lipid species co-enriched in circulating EVs and EVs from cell culture. A.** Scatter plot showing relative abundance of lipids in pEVs vs p100K and pEVs vs pDGS.LD. Lipids in blue are enriched in pEVs. **B.** Heatmap depicting differentially abundant lipids in pEVs vs NonEVs. **C.** Scatter plot showing relative abundance of lipids in pEVs vs p100K/pDGS.LD and in vitro EVs vs WCL lipidome data sets. Blue circles (lipid markers) represent lipids with significantly greater abundance (Log2 FC > 0.5 & p value < 0.05) in pEVs vs pDGS.LD and in vitro EVs vs cells. Red circles (exclusion lipids) represent lipids with significantly lower abundance (Log2 FC < -0.5 & p value < 0.05) in pEVs vs pDGS.LD and in vitro EVs vs cells. Grey circles represent lipids that do not meet the above criteria. **D-E.** Bar plots showing relative abundance (log2 centred intensities) of differentially abundant (p<0.05) lipids in indicated lipidomes.

**Supplementary** Figure 11**. PS lipid class is enriched in EVs in plasma. A** Lipid abundance (normalized) across 12 fractions of density gradient separated plasma. **B.** Box plots depict abundance of cluster 1-4 lipids for DGS fractions for each biological replicate (R1-3). Y axis represents Z-scored abundance (MS-based abundance for each lipid - mean abundance) / standard deviation (Z score normalization). Blue lines highlights fractions 6-7 (corresponding to pEV fractions), whereas red line depicts fractions 1-3 (corresponding to pDGS.LD fractions).

**Supplementary** Figure 12**. Lipidome analysis of pEVs from AusDiab validation cohort and non-transformed cells**. **A.** Boxplot depicting normalised abundance (vsn normalized) of lipids quantified in non-transformed EVs (in vitro Small EVs or in vitro Large EVs) or pEVs, NonEVs or neat plasma from AusDiab validation set. **B.** Heatmap of Pearson correlation of quantified lipids.

**Supplementary Figure 13. Heatmap of lipid features distribution in pEVs versus NonEVs from AusDiab validation cohort and EVs from non-transformed cells.**

**Supplementary** Figure 14**. Machine learning classification of EV and NonEV particles in plasma using protein features. A.** Confusion matrix (using the ensemble model) of training set (70%), remaining validation set (30%), and independent test set, for cluster 1 proteins, which include top 97 proteins enriched in EVs (FC>5) and 28 exclusion proteins. Probability scores for sample classification into EVs using ensemble model on independent test set. Greater scores indicate higher confidence in predicting sample belonging to EV class. **B.** Confusion matrix using nnet classifier of validation set (30%) and independent test set, for 126 pEV surface proteins. Bar plot show probability scores for sample classification into EVs using nnet model on independent test set. **C.** Confusion matrix using nnet classifier of validation set (30%) and independent test set, for ADAM10 vs sample median intensities for each proteome as depicted in Figure 6B.

**Supplementary** Figure 15**. Machine learning classification of EV and NonEV particles in plasma using lipid features. A** Confusion matrix using Neural Network algorithm (’nnet’) classifier of training set (70%), validation set (30%) and independent test set, for cluster c1 and c4 lipids (upper panel), and top 20 c1 and c4 lipid features following reclusive feature elimination (lower panel). **B.** Bar plot indicated probability scores for DGS fraction lipidomes classification into EVs using ’nnet’ model based on top 20 c1 and c4 lipid features on independent test set. Greater scores indicate higher confidence in predicting DGS fractions belonging to EV class.

**Supplementary** Figure 16**. Spectral flow cytometry (Cytek Aurora)-based pEV analysis for ADAM10 and PS expression A.** Scatter plot depicts instrument gating calibration using 90 nm, 125 nm, 150 nm, 200 nm and equal mix (90-200 nm) beads. Bottom panel indicated buffer alone control. YG3-A channel used for detecting Alex Flour 568 signal. R1-A channel used for detecting APC signal. **B.** Cytek Aurora-based pEV analysis. Scatter plot showing detection of ADAM10 antibody-AF568 positive (detected using YG3-A channel) and PS positive (Annexin V-APC, detected using R1-A channel) in pEVs. IgG used as control for ADAM10 antibody. SDS detergent (0.5%) solubilization was performed to indicate EV origin of these signals. **C.** Bar plot depicting ADAM10+ (AF548 signal), PS+ (APC signal) or overlapping (ADAM10 and PS +) signal.

**Supplementary Figure 17. Interrogating protein of interest expression in circulating EVs using the Shiny app.**

**Supplementary** Figure 18. **Interrogating lipid of interest expression in circulating EVs using the Shiny app.**

Supplementary Figure 19. Characterization of PS 36:1 in EVs using negative ionization mode. Pooled EV’s were run under the same conditions in negative ionization mode, looking for fragments corresponding to the neutral loss of the serine headgroup (red) and all possible fatty acid species, resulting in identification of 18:0 (283.3 *m/z*) and 18:1 (281.3 *m/z*).

Supplementary Figure 20. Validation of isobars and isomer separation between PC 35:2, PC O-36:2 and PC P-36:1 with chromatography. A - Transition (772.6/184.1) corresponding to PC 35:2 / PC O-36:2 / PC P-36:1 using pooled human plasma (0.1μl on column injection) run on an Agilent 6495C with chromatographic conditions as previously described (Agilent Infinity II HPLC). B - EIC’s corresponding to PC 35:2 and PC O-36:2 / PC P-36:1 using pooled human plasma (0.1μl on column injection) run on an Agilent 6546 QTOF with chromatographic conditions as previously described. Resolution is approximately 60,000 FWHM. The *m/z* 772.5851 (C43H83NO8P) putatively corresponds to PC 35:2 (Top panel) while 772.6215 (C44H87NO7P) corresponds to PC O-36:2 or PC P-36:1 (Panel). C,D - EIC’s corresponding to PC 35:2 and PC O-36:2 / PC P-36:1 using pooled human plasma (0.1μl on column injection) run on an HFx Orbitrap and a Vanquish analytical HPLC with chromatographic conditions as previously described. Analysis was run as full MS1 scan with resolution set to 240,000 FWHM. The EIC 772.6215 (C44H87NO7P) corresponds to PC O- 36:2 or PC P-36:1 (top panel) while EIC 772.5851 (C43H83NO8P) putatively corresponds to PC 35:2 (bottom panel). E - Lithium acetate was added to the running solvent. Species corresponding to PC 35:2 was measured as their lithium adduct [M+Li]^+^ m/z 778.6, with the loss of the phosphocholine headgroup as the product ion (m/z 595.2). Product ions corresponding to the acyl composition 17:0 (449.2) and 18:2 was observed (439.2) split across two distinct chromatographic peaks. This was then followed up by subsequent synthesis of branched and straight isomer standards^134^. F - Transition (772.6/184.1) corresponding to PC 35:2 / PC O-36:2 / PC P-36:1 using pooled human plasma (0.1μl on column injection) run on an Agilent 6490 with chromatographic conditions as previously described. Red trace, HCl vapor treated, Blue trace, untreated control.

**Supplementary Figure 21. Uncropped Western blot images.**

---

**Extended data 1.**
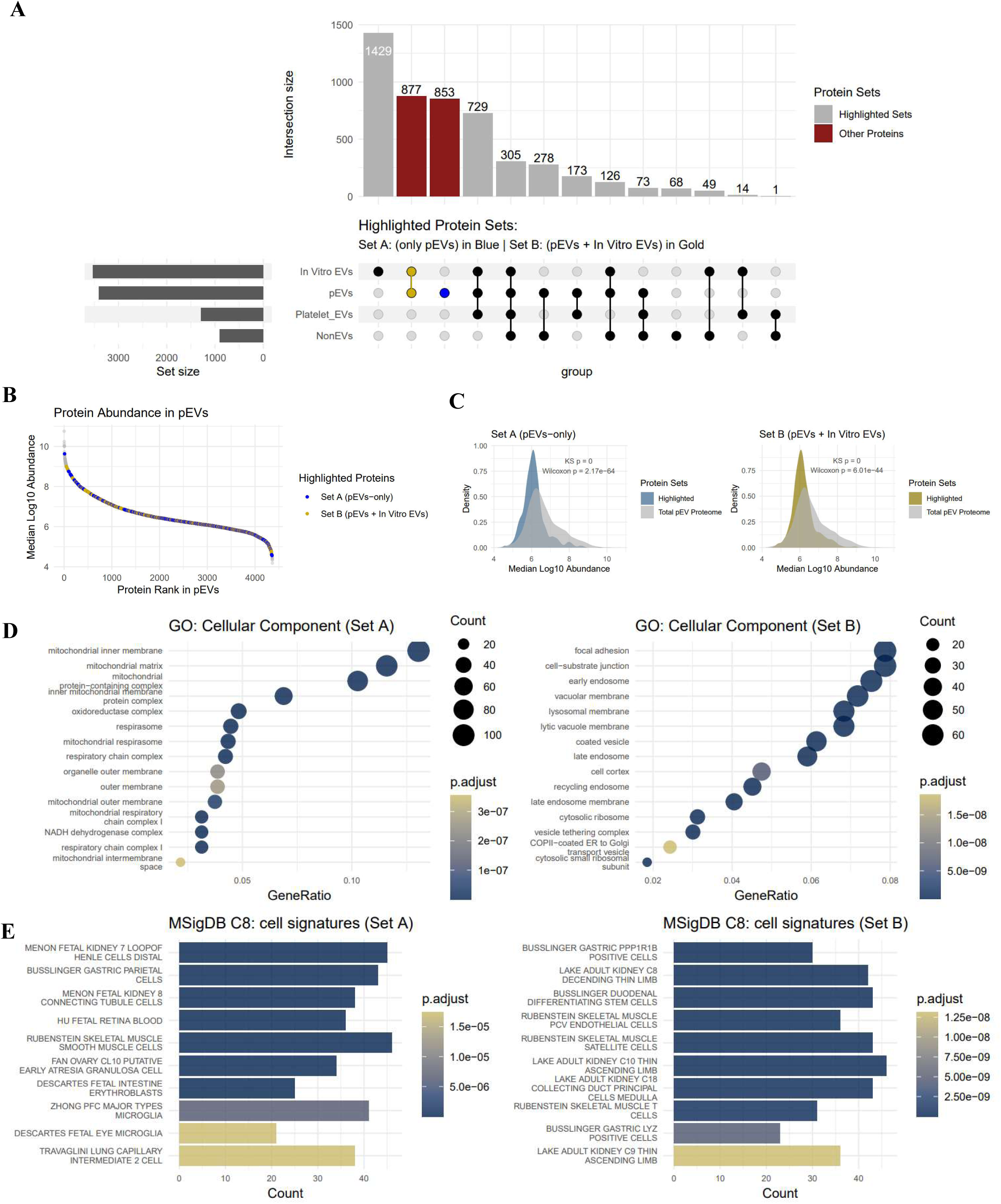
Diverse cellular source attribution to pEV proteome. **A**. Distribution of activated platelet EV proteins in pEVs and in vitro EVs. Set A and Set B proteins represent proteins quantified exclusively in pEVs and in vitro EVs versus activated platelet EVs^53^. **B.** Distribution of Set A and Set B proteins (abundance) in pEV proteome. **C.** Density plots of Set A and Set B protein abundance vs pEV proteome abundance. **D.** Gene Ontology (Cellular Component) analysis of Set A and Set B proteins. **E.** Gene Set Enrichment Analysis (using Molecular signature database^133^ for cell signatures (C8 category)) of Set A and Set B proteins.

**Extended data 2.**
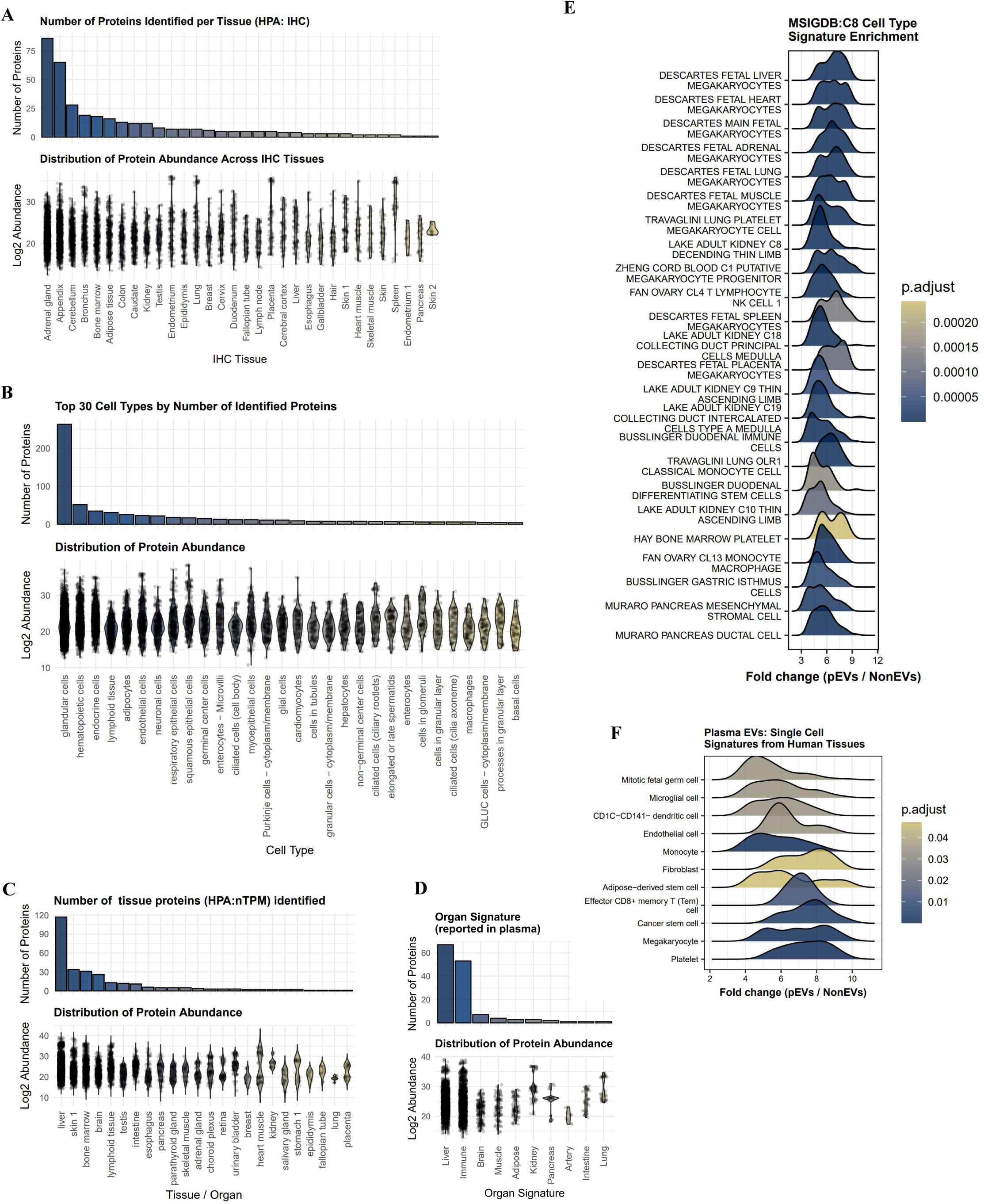
Quantification of tissue / organ-associated proteins in pEVs. **A**. Number of tissue-associated proteins (based on Human Protein Atlas (HPA) immunohistochemistry data (IHC)^56^) quantified in pEV proteome. Abundance distribution of proteins are presented in lower panel. **B.** Number of cell-type-associated proteins (based on HPA IHC) quantified in pEV proteome. Abundance distribution of proteins is presented in lower panel. **C.** Number of tissue-associated proteins (based on HPA normalized transcript per million (nTPM)) quantified in pEV proteome. Abundance distribution of proteins is presented in lower panel. **D.** Number of organ-associated proteins reported in plasma using aptamers quantified in pEV proteome^57^. Abundance distribution of these proteins is presented in lower panel. **E.** Gene Set Enrichment Analysis (using Molecular signature database^133^ for cell signatures (C8 category) of proteins enriched in pEV vs NonEV proteomes. **F.** Enrichment of cell signatures based on single-cell sequencing studies of human tissue^133^ in proteins enriched pEV vs NonEVs.

**Extended data 3.**
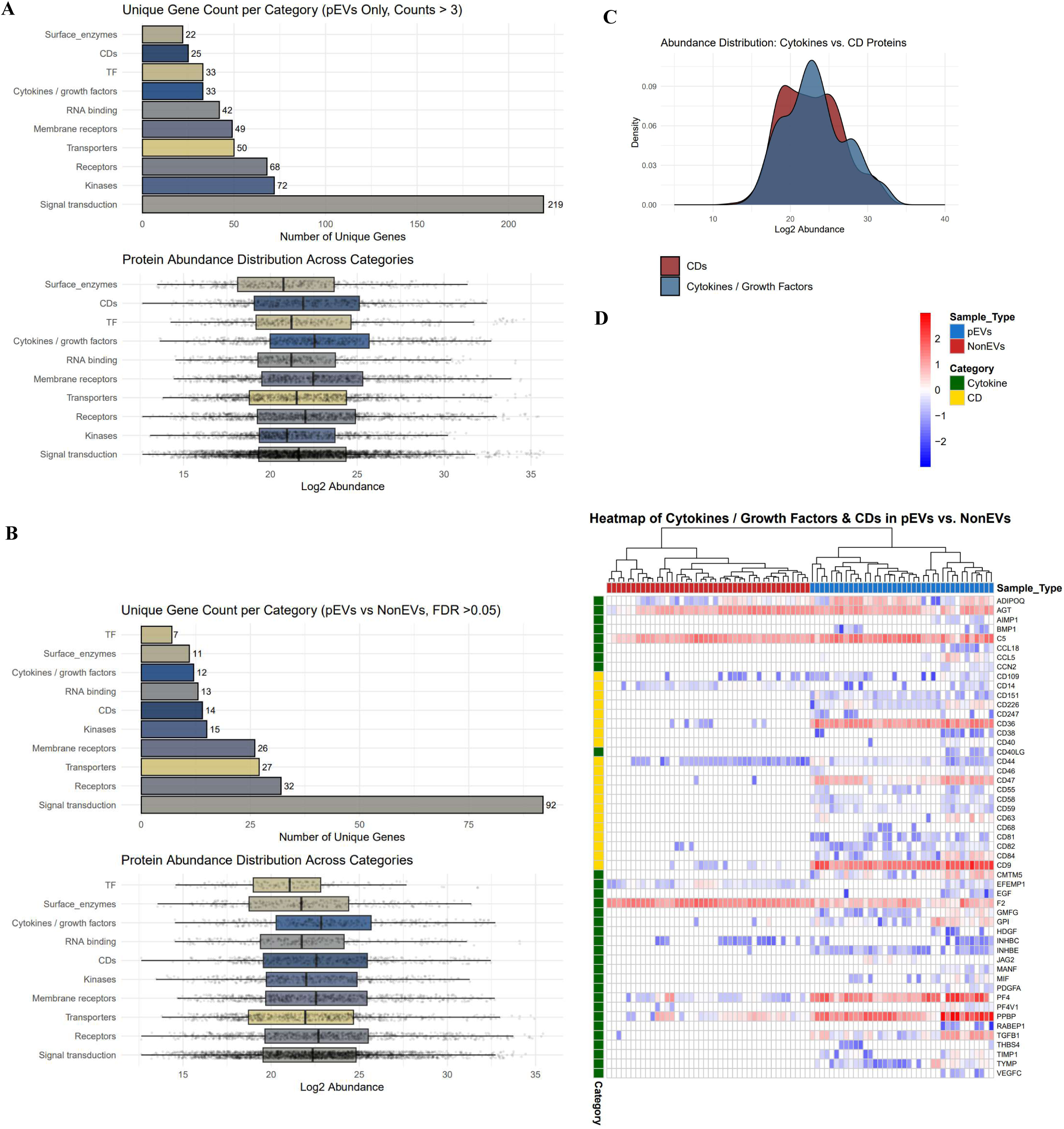
Diversity of bioactive cargo in pEVs. **A**. Number of different cargo-type quantified in pEV proteomes (in at least 3 out of 42 pEV proteomes), with box-plot in lower panel representing protein abundance distribution. TF, transcription factors; CDs, Cluster of Differentiation. Gene Ontology terms related to cytokine activity (GO:0005125), chemokine activity (GO:0008009), growth factor activity (GO:0008083), and signal transduction (GO:0007165) to ensure comprehensive coverage of bioactive EV-associated molecules **B.** Number of different cargo-type quantified in pEV proteomes (but significantly enriched in pEVs vs NonEVs, FDR <0.05), with box-plot in lower panel representing protein abundance distribution. **C.** Abundance distribution of cytokines/growth factors and CD proteins quantified in pEV proteome. **D.** Heatmap of cytokines and CD proteins quantified in pEV and NonEV proteomes. Grey boxes in the heatmap represent non-quantification.

**Extended data 4.**
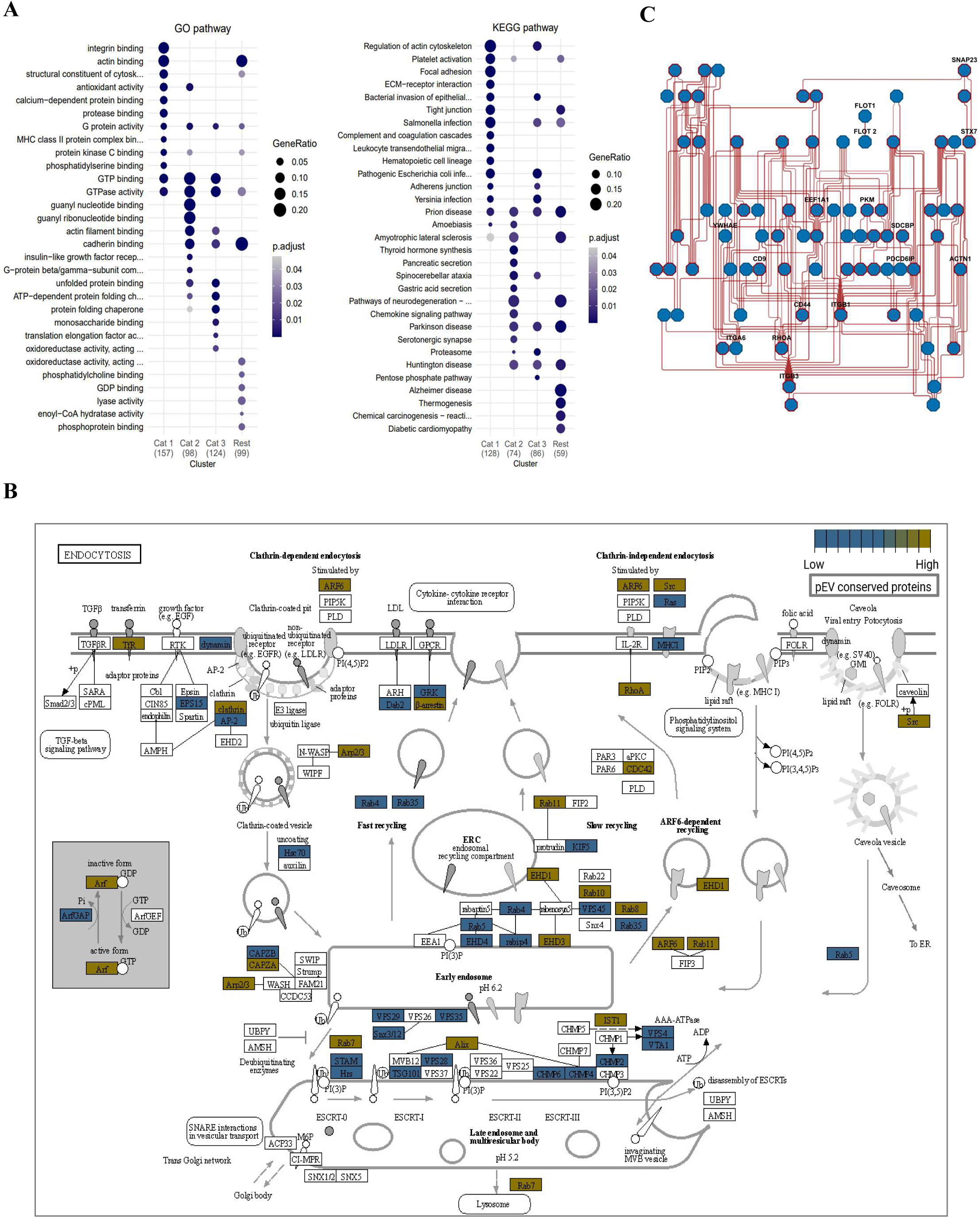
Characterization of conserved proteins in pEVs. **A**. Gene Ontology pathways and KEGG pathways enriched (p<0.05) in each cluster. **B.** KEGG endocytosis pathway graph rendered by Pathview where category 1 proteins are indicated as high and category2-3 proteins are indicated as low. **C.** Protein–protein interaction networks of cluster 1 proteins analysed using STRING-DB and Cytoscape. Each node represents a protein. Nodes with red border are enriched for “vesicular pathway”.

**Extended data 5.**
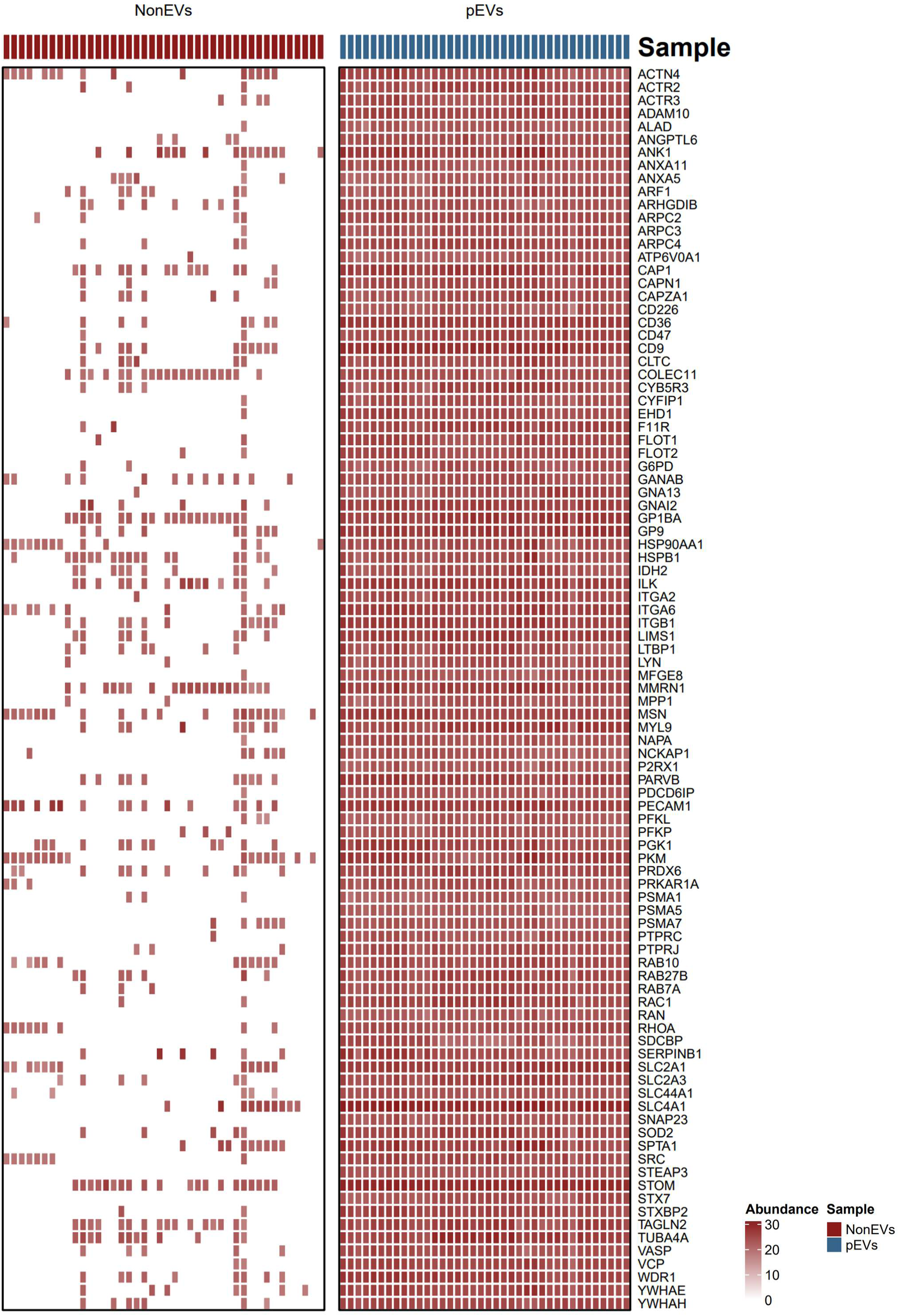
Heatmap of selected pEV protein features.

**Extended data 6.**
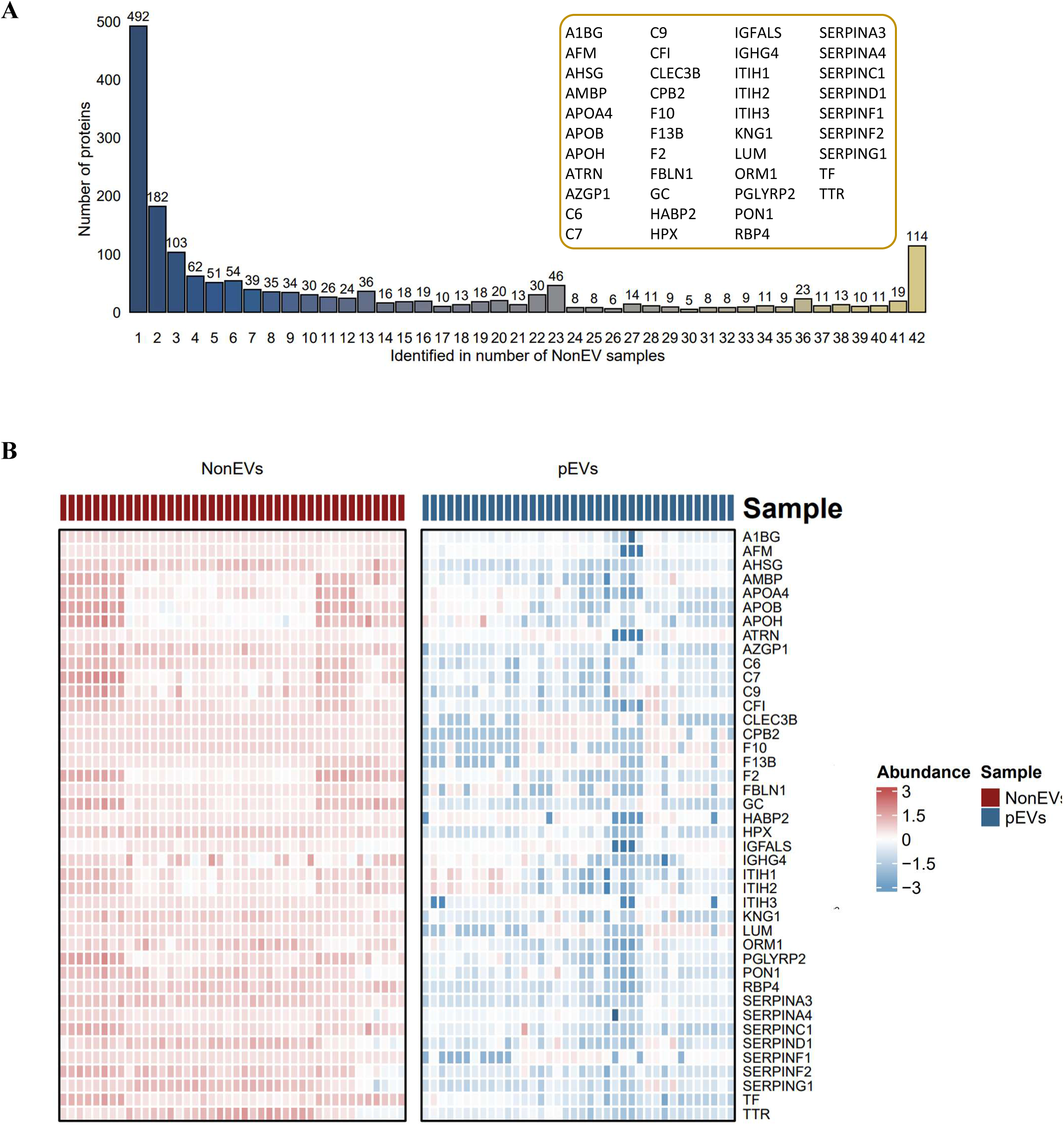
Mapping the core proteome of NonEVs in humans. **A.** Occurrence analysis of proteins in 38 pEV proteomes in our data set where Category 1 proteins are ubiquitously quantified. **B.** Heatmap of conserved protein features in NonEVs.

**Extended data 7.**
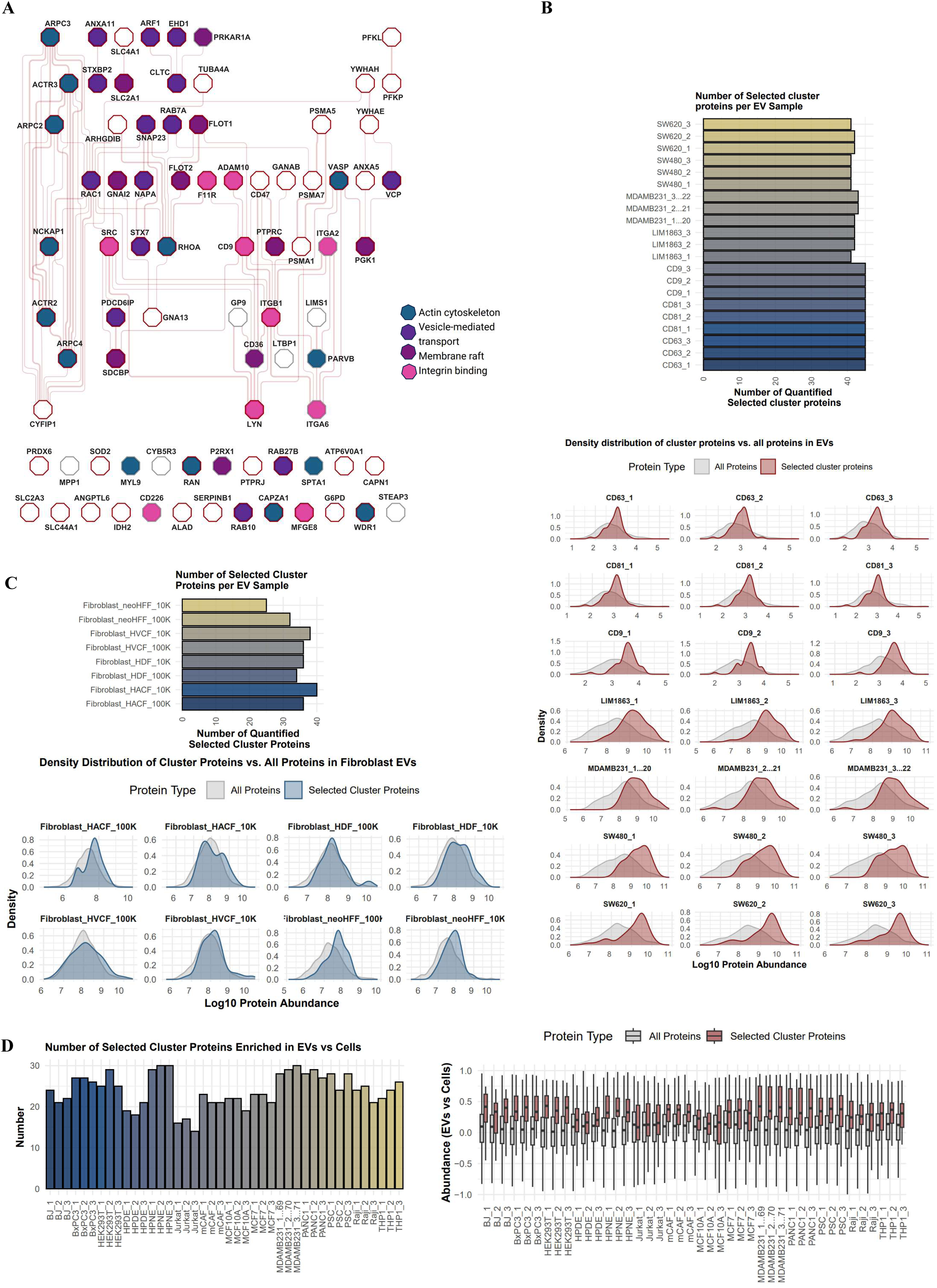
Coordinated molecular patterns across EV proteomes. **A.** Protein– protein interaction networks of selected EV protein features (quantified in <30% of NonEV datasets) analyzed using STRING-DB and Cytoscape; each node represents a protein. Functional enrichment (GO cellular component) was performed using Cytoscape. Nodes with red border are enriched for “Extracellular exosome” term. **B**. Number of proteins belonging to coordinated molecular patterns (in Panel A) quantified in in vitro EVs (n=3 per cell line) or EV subtypes (CD81+, CD63+, and CD9+ EVs) prevalent in the circulatory system^70^. Quantification in MS data of individual biological replicates are shown. Lower panel shows distribution of protein abundance of these molecular features against all other proteins in their respective proteome. **C.** Number of proteins belonging to coordinated molecular patterns (in Panel A) quantified in small EVs (100,000 g ultracentrifuged EVs 100K) or large EVs (10,000 g ultracentrifuged EVs 10K) from non-transformed cells (fibroblasts). Lower panel shows distribution of protein abundance of these molecular features against all other proteins in their respective proteome. **D.** Number of proteins belonging to coordinated molecular patterns (in Panel A) quantified in small EVs from 14 cell lines as reported^30^. Lower panel shows relative abundance of these molecular features in EVs versus their parental cell proteomes.

**Extended data 8.**
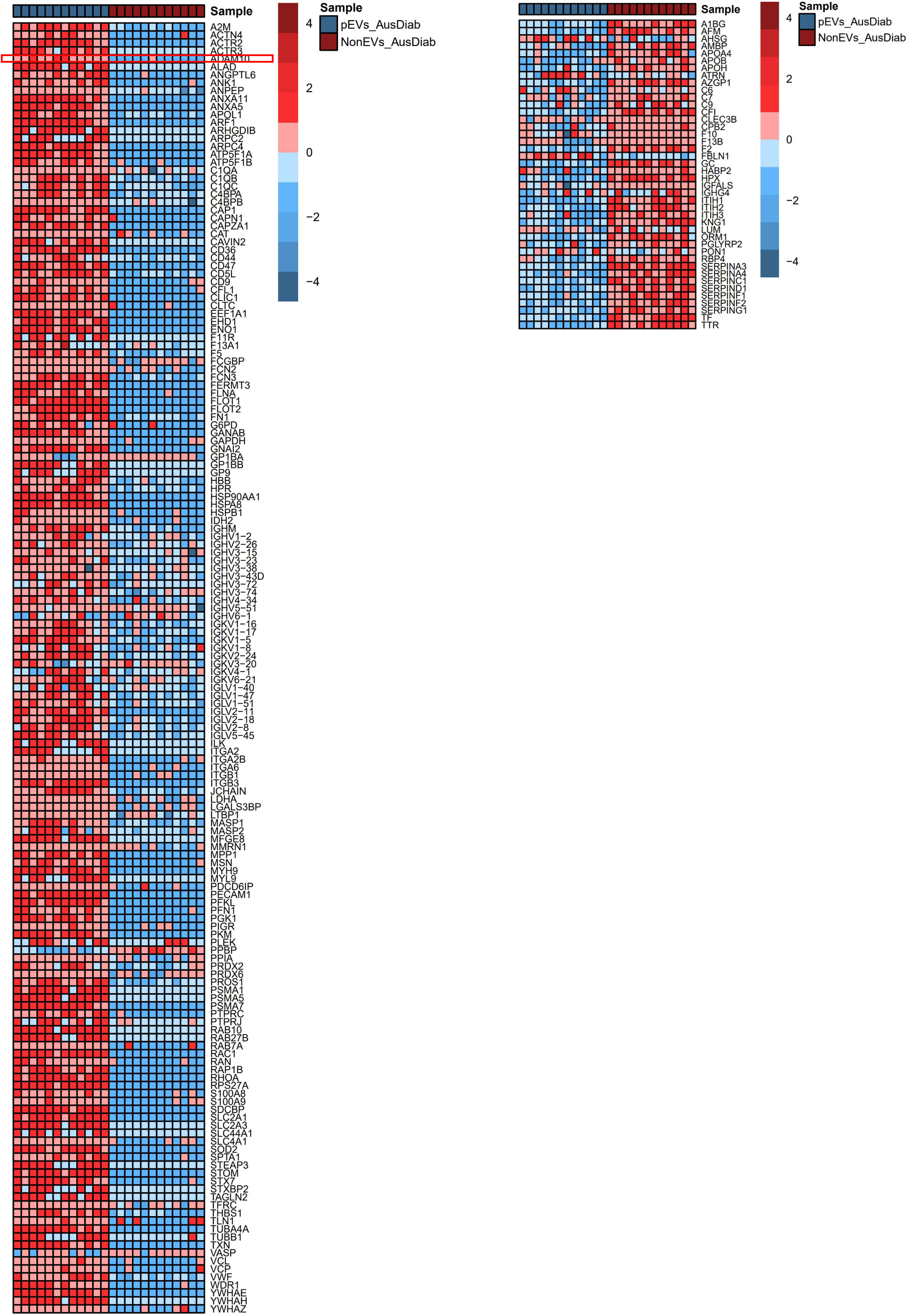
Heatmap of pEV and NonEV protein features in AusDiab validation set.

**Extended data 9.**
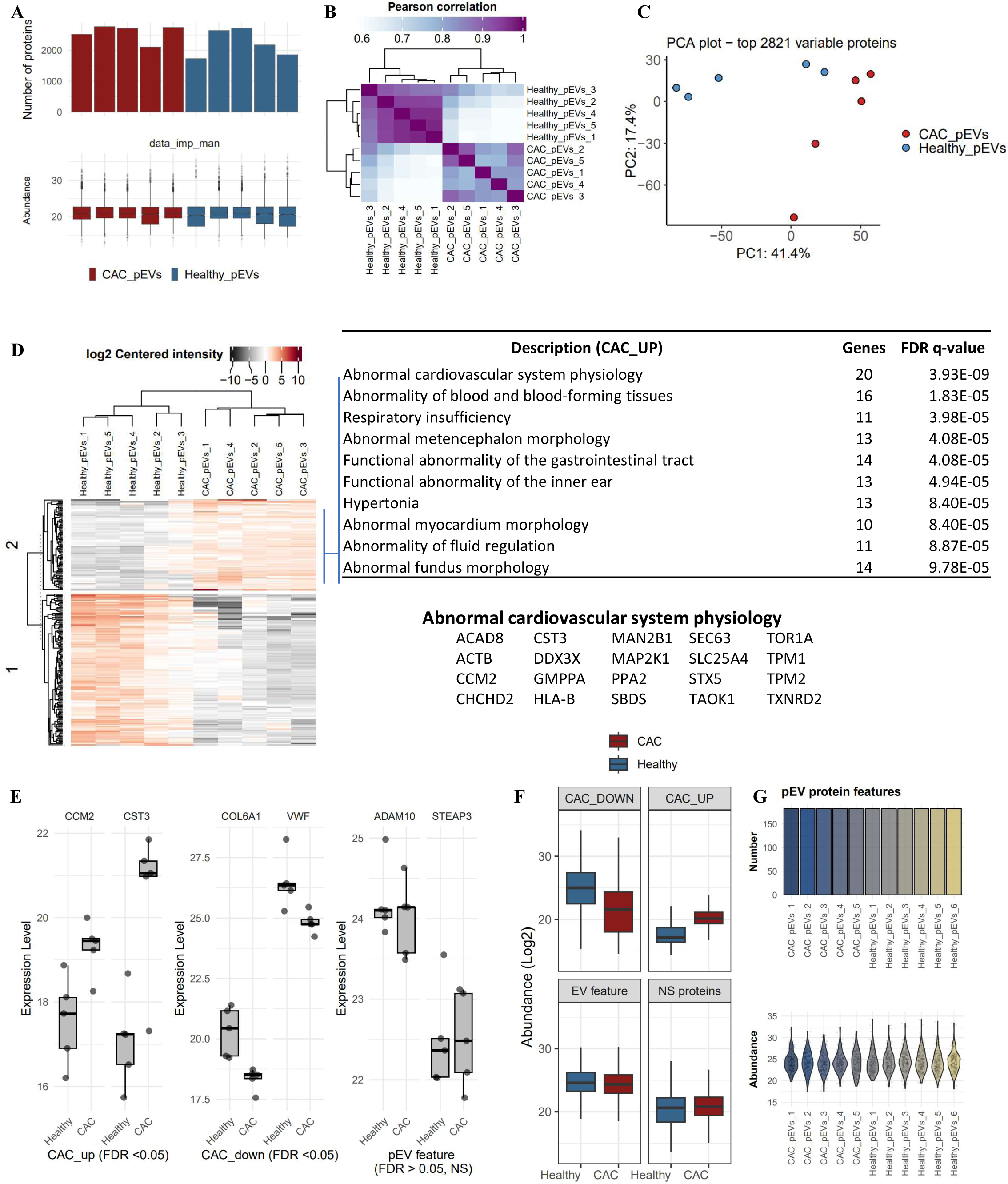
Proteome analysis of pEVs from individuals with either positive CAC score (CAC group) or zero CAC score (Healthy group). **A**. Proteins quantified in CAC_pEVs or Healthy_pEVs. Lower panel indicates normalized proteins abundance (vsn-normalized) **B.** Pearson correlation of quantified proteins. **C.** Principal component analysis of quantified proteins. **D.** Heatmap depicting K-means clustering of differentially abundant proteins (FDR <0.05) in CAC_pEVs versus Healthy_pEVs: 76 upregulated and 127 downregulated proteins (FDR <0.05), with GO pathways enriched in upregulated proteins. Proteins of the GO pathway ‘Abnormal cardiovascular system physiology’ are shown. **E.** Boxplot showing vsn-normalizsed abundance (log2) of indicated proteins in CAC_pEVs and Healthy_pEVs proteomes. **F.** Boxplot showing distribution of either significantly upregulated (CAC_UP) or down regulated (CAC_DOWN) proteins, or non-significantly dysregulated proteins (NS proteins), and EV proteins features (EV feature) in CAC_pEVs and Healthy_pEVs proteomes. **G.** pEV protein features quantified in individual CAC_pEVs versus Healthy_pEVs proteomes. Lower panel shows distribution of pEV protein features in individual proteomes.

**Extended data 10.**
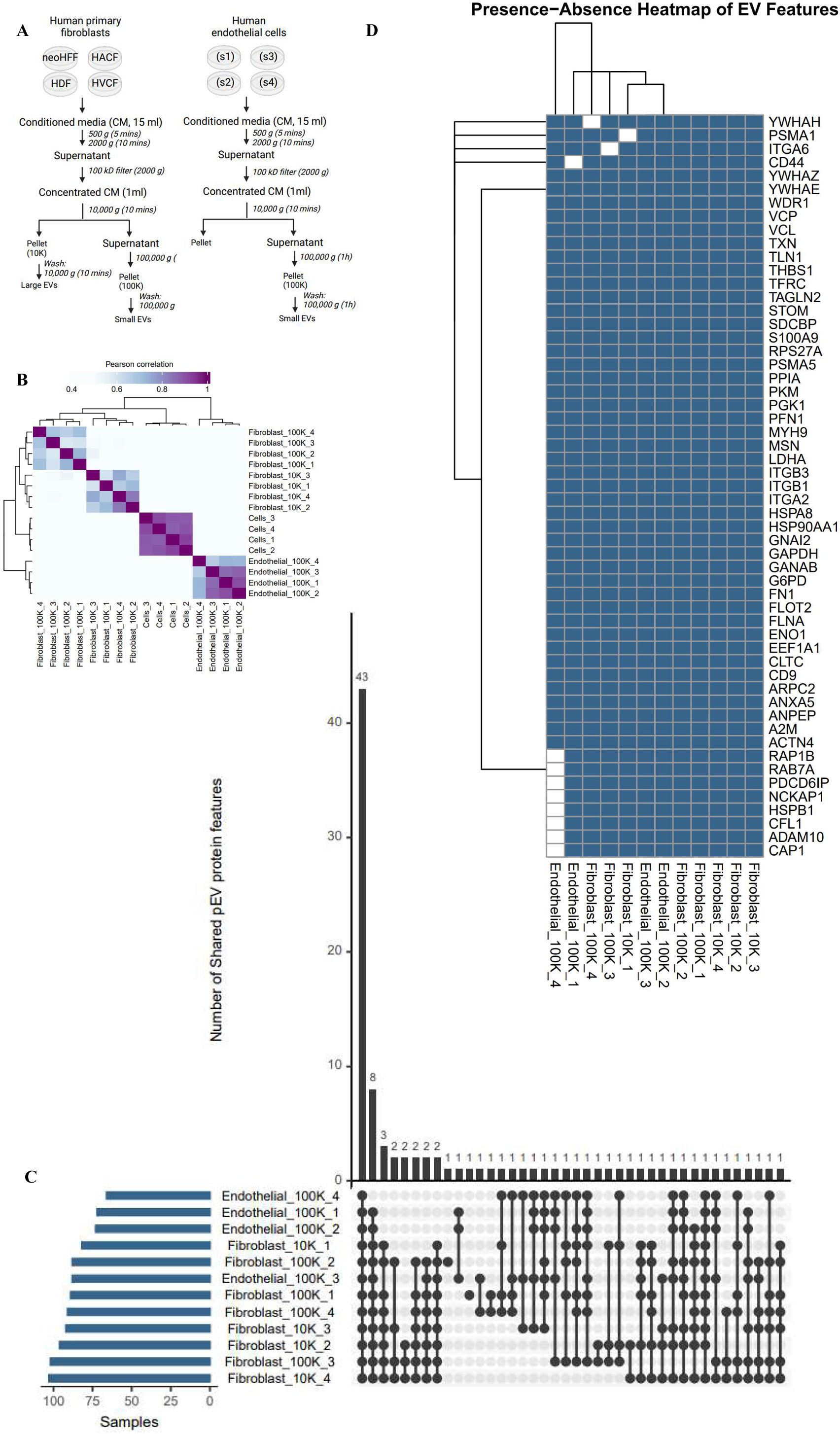
Conservation of pEV protein features in EVs from non-transformed cells. **A**. Workflow for obtaining small EVs (100K) or large EVs (10K) from conditioned media of either human primary fibroblasts or human primary endothelial cells. **B.** Pearson correlation of quantified proteins in indicated proteomes using mass spectrometry. Fibroblasts (whole cell lysates) were used as reference proteome for cells. **C.** pEV protein features quantified in small EVs (100K) or large EVs (10K) released by fibroblasts or endothelial cells. D. Heatmap indicates detection of pEV protein features in small EVs (100K) or large EVs (10K) released by fibroblasts or endothelial cells.

**Extended data 11.**
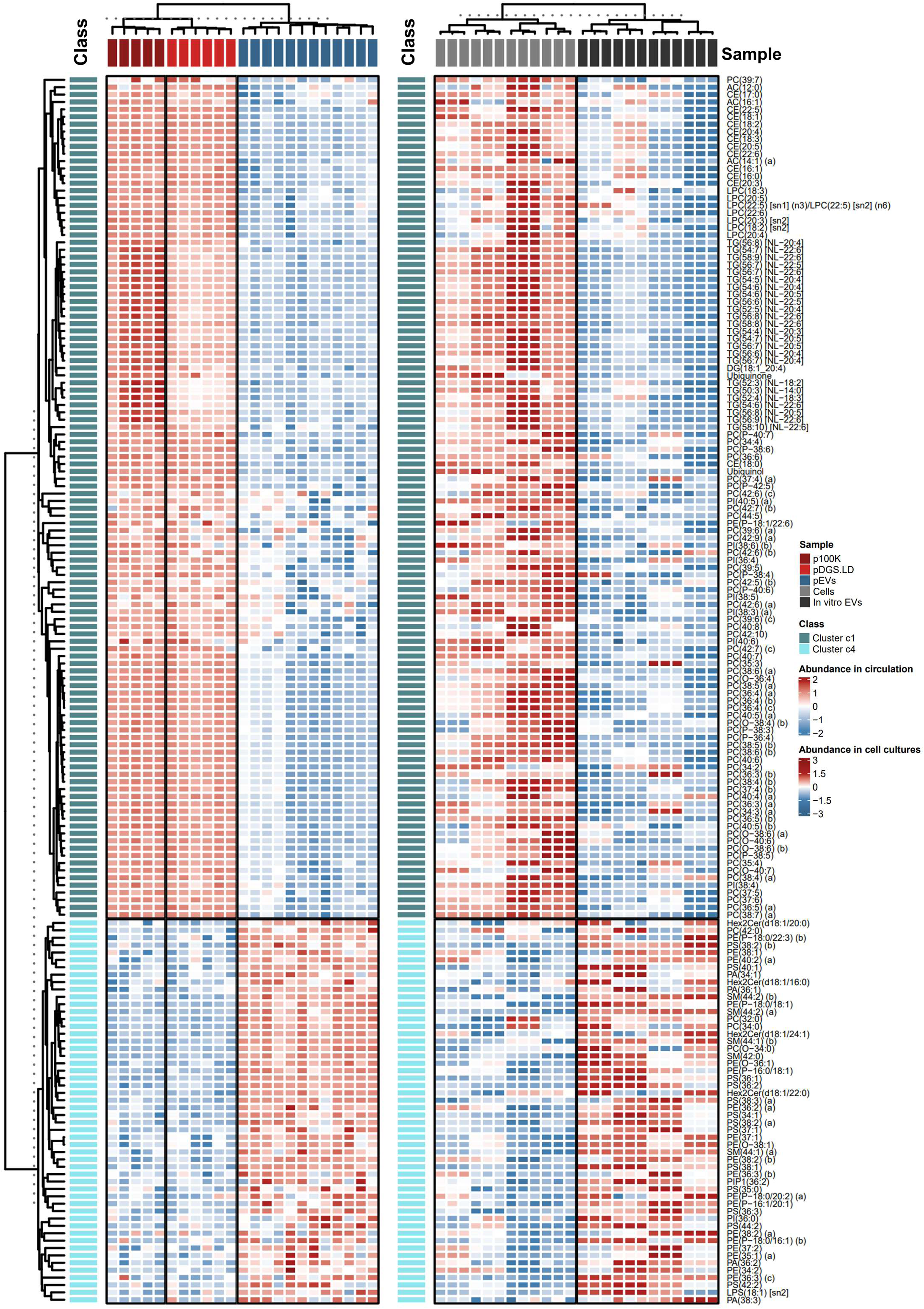
Heatmap of EV cluster 1 and cluster 4 lipids in EVs and NonEVs.

**Extended data 12.**
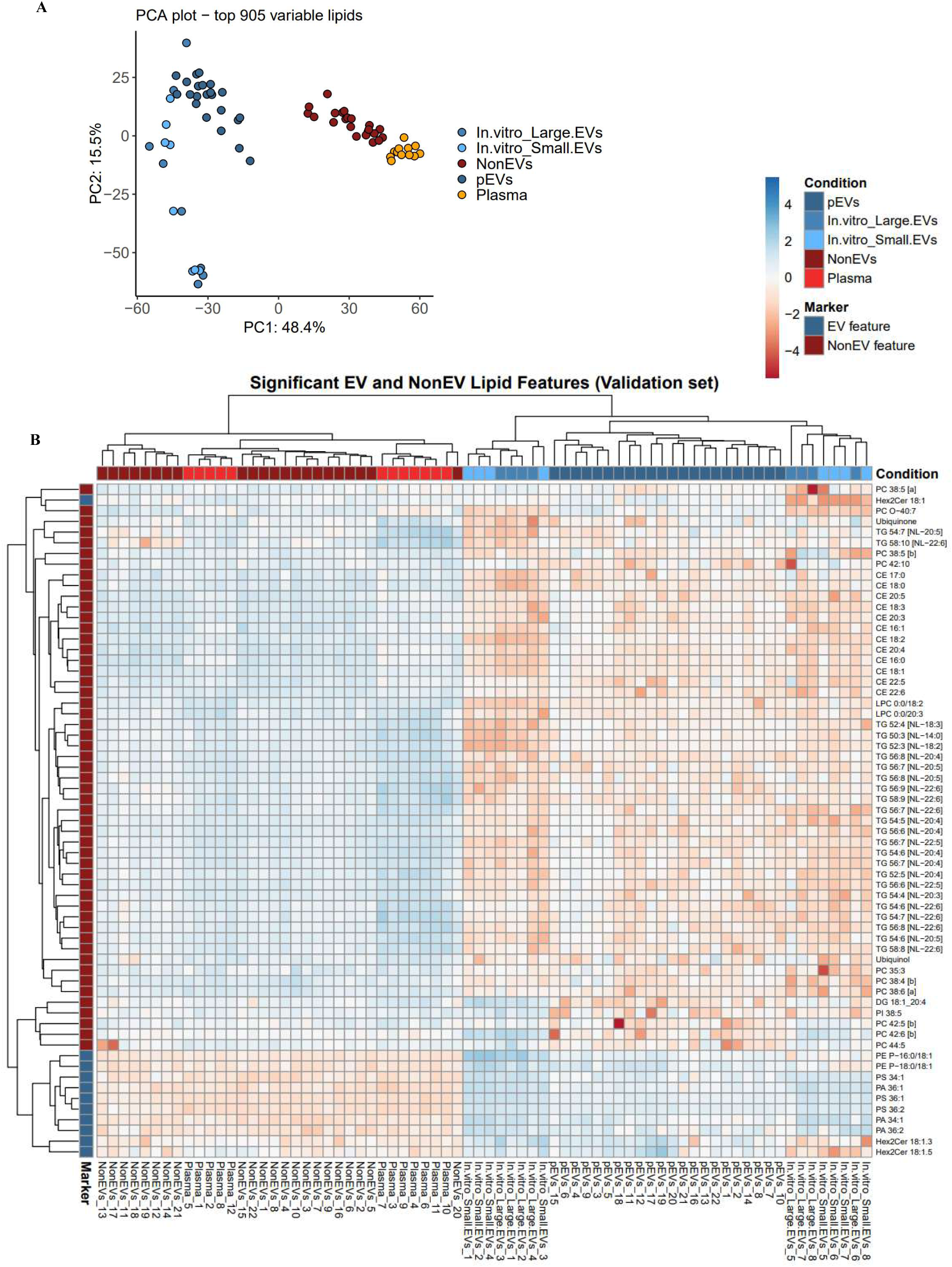
Conservation of pEV lipid features in EVs from AusDiab validation set and in non-transformed cells. **A**. Principal component analysis of quantified lipids in non-transformed EVs (in vitro Small EVs or in vitro Large EVs) or pEVs, NonEVs or neat plasma from AusDiab validation set. **B.** Heatmap depicts distribution of pEV and NonEV lipid features quantified in AusDiab validation set (pEVs, NonEVs or neat plasma) and in non-transformed cell-derived EVs (in vitro Small EVs or in vitro Large EVs).

**Extended data 13.**
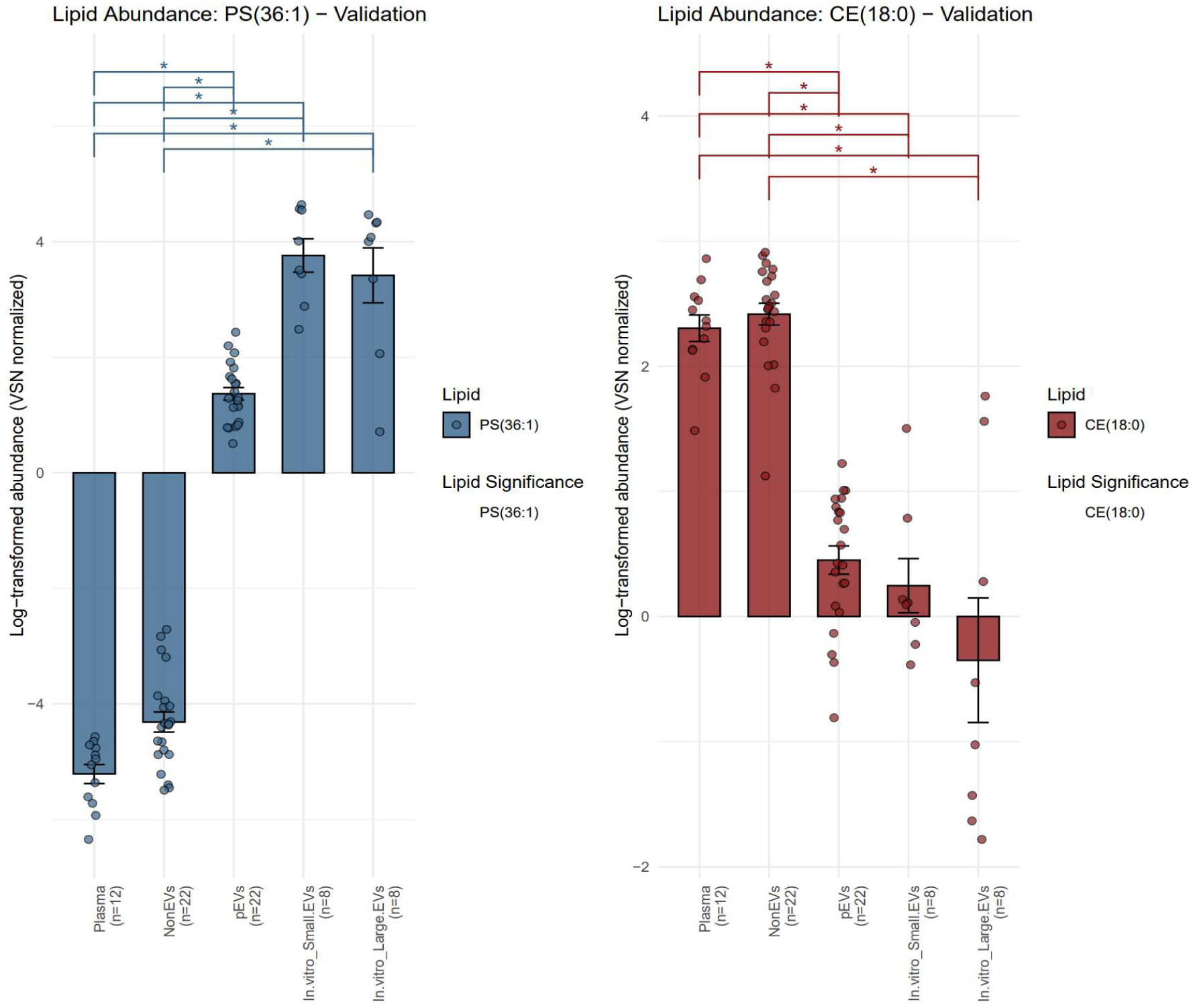
Relative abundance of pEV lipid features PS(36:1) and CE(18:0) in pEVs lipidome datasets. Lipid features in AusDiab validation set (pEVs, NonEVs or neat plasma) and in non-transformed cell-derived EVs (in vitro Small EVs or in vitro Large EVs).

